# Harnessing machine learning to predict antibiotic susceptibility in *Pseudomonas aeruginosa* biofilms

**DOI:** 10.1101/2025.04.24.650389

**Authors:** Fauve Vergauwe, Gaetan De Waele, Andrea Sass, Callum Highmore, Niall Hanrahan, Yoshiki Cook, Mads Lichtenberg, Margo Cnockaert, Peter Vandamme, Sumeet Mahajan, Jeremy S. Webb, Filip Van Nieuwerburgh, Thomas Bjarnsholt, Willem Waegeman, Tom Coenye

## Abstract

Standard antibiotic susceptibility tests (ASTs) often fail to accurately predict treatment outcomes because they do not account for biofilm-specific mechanisms of reduced susceptibility. In the present study, we explored alternative approaches to predict tobramycin susceptibility of *Pseudomonas aeruginosa* biofilms that were experimentally evolved in physiologically relevant conditions. To this end, we used four analytical methods – whole-genome sequencing (WGS), matrix-assisted laser desorption/ionization-time of flight mass spectrometry (MALDI-TOF MS), isothermal microcalorimetry (IMC) and multi-excitation Raman spectroscopy (MX-Raman). Machine learning models were trained on data outputs from these methods to predict tobramycin susceptibility of our evolved strains and subsequently validated with a collection of clinical isolates. For minimal inhibitory concentration (MIC) predictions of the evolved strains, the highest accuracy^±1^ was achieved with MALDI-TOF MS (97.83 %), while for biofilm prevention concentration (BPC) predictions, Raman spectroscopy performed best with an accuracy^±1^ of 80.43 %. Overall, all analytical methods demonstrated comparable predictive performance, showing their potential for improving biofilm AST.

## Introduction

Antibiotic susceptibility tests (ASTs) guide the selection of antimicrobial agents for treating infections ^1,2^. These ASTs typically evaluate the response to antibiotics of planktonic bacteria in suspension (e.g. broth microdilution test ^3^) or bacteria grown on agar surfaces (e.g. disk diffusion test ^4^). However, the conditions of these *in vitro* tests do not represent the *in vivo* microenvironment, where bacteria often occur as biofilms, i.e. aggregates encased in extracellular matrix derived from either the bacteria or the host ^5^. Biofilm-growing bacteria exhibit distinct physiological characteristics that confer increased tolerance and resistance to antibiotics compared to their planktonic counterparts ^6,7^ and they can withstand antibiotic concentrations up to 100-1,000 times higher than those effective against planktonic cells ^8–11^. As a result, conventional ASTs frequently fail to predict treatment success, as the biofilm phenotype is not considered ^8,12,13^. To improve *in vitro* ASTs, culture media that more closely mimic the *in vivo* microenvironment can be used. An example of such a medium is synthetic cystic fibrosis medium 2 (SCFM2), which simulates the lung environment of cystic fibrosis (CF) patients ^14^. In SCFM2, *Pseudomonas aeruginosa* forms biofilm microaggregates that closely resemble those found in the sputum of CF patients ^15^ and exhibit similar gene expression profiles ^16^, and this medium can be used to determine biofilm susceptibility ^10^.

However, biofilm-based ASTs remain challenging to implement and are time consuming. In the present study we explored alternative methods to predict the antibiotic susceptibility of *P. aeruginosa* biofilms, using whole-genome sequencing (WGS), matrix-assisted laser desorption/ionization-time of flight mass spectrometry (MALDI-TOF MS), isothermal microcalorimetry (IMC) and multi-excitation Raman spectroscopy (MX-Raman). Each of these analytical techniques provides unique insights into different bacterial properties. WGS allows the prediction of antibiotic susceptibility by confirming the presence of known resistance genes or mutations ^17–19^; however, it may be less suitable to detect poorly understood and/or novel tolerance and resistance mechanisms. MALDI-TOF MS, widely used in clinical microbiology laboratories for rapid pathogen identification, also holds potential for antibiotic susceptibility testing ^20–22^. Machine learning algorithms trained on large MALDI-TOF MS datasets have been successfully used to classify isolates as susceptible or resistant ^23^. IMC measures heat production associated with metabolic processes. Microbial metabolism plays an important role in antimicrobial susceptibility ^6,24^, and measuring metabolic activity with IMC can be used to evaluate the activity of antibiotics ^25–27^. Lastly, Raman spectroscopy (RS) measures the inelastic scattering of light due to molecular vibrations, thereby offering information about the molecular composition of a sample ^28^. In MX-Raman, multiple wavelengths are used to excite the sample, providing a more comprehensive fingerprint of bacterial cells ^29^. Several studies have demonstrated the potential of RS to differentiate between resistant and susceptible bacteria ^30,31^. Using WGS, MALDI-TOF MS, IMC and MX-Raman data obtained from experimentally evolved *P. aeruginosa* strains, we developed an ordinal regression model to predict the minimal inhibitory concentration (MIC, as a measure of susceptibility of planktonic cells) and the biofilm prevention concentration (BPC) of tobramycin. The BPC was defined as the lowest concentration of antimicrobial agent that prevented at least 90 % of biofilm growth compared to the growth control in SCFM2 ^13^. For MIC predictions, all techniques achieved an accuracy^±1^ > 89 % (allowing a margin of error of one dilution), with MALDI-TOF MS achieving the highest performance by predicting the MIC with an accuracy^±1^ of 97.83 %. For BPC predictions, all techniques showed an accuracy^±1^ > 73 %; the best BPC predictions were obtained with Raman spectroscopy at 532 nm, achieving an accuracy^±1^ of 80.43 %. After training, the model was validated with a set of *P. aeruginosa* isolates recovered from CF patients. For tobramycin MIC predictions, IMC and MALDI-TOF MS scored above random predictions, with IMC showing the best performance. For BPC predictions, only IMC performed above random predictions. An overview of the workflow is presented in Fig. 1.

**Figure 1.**
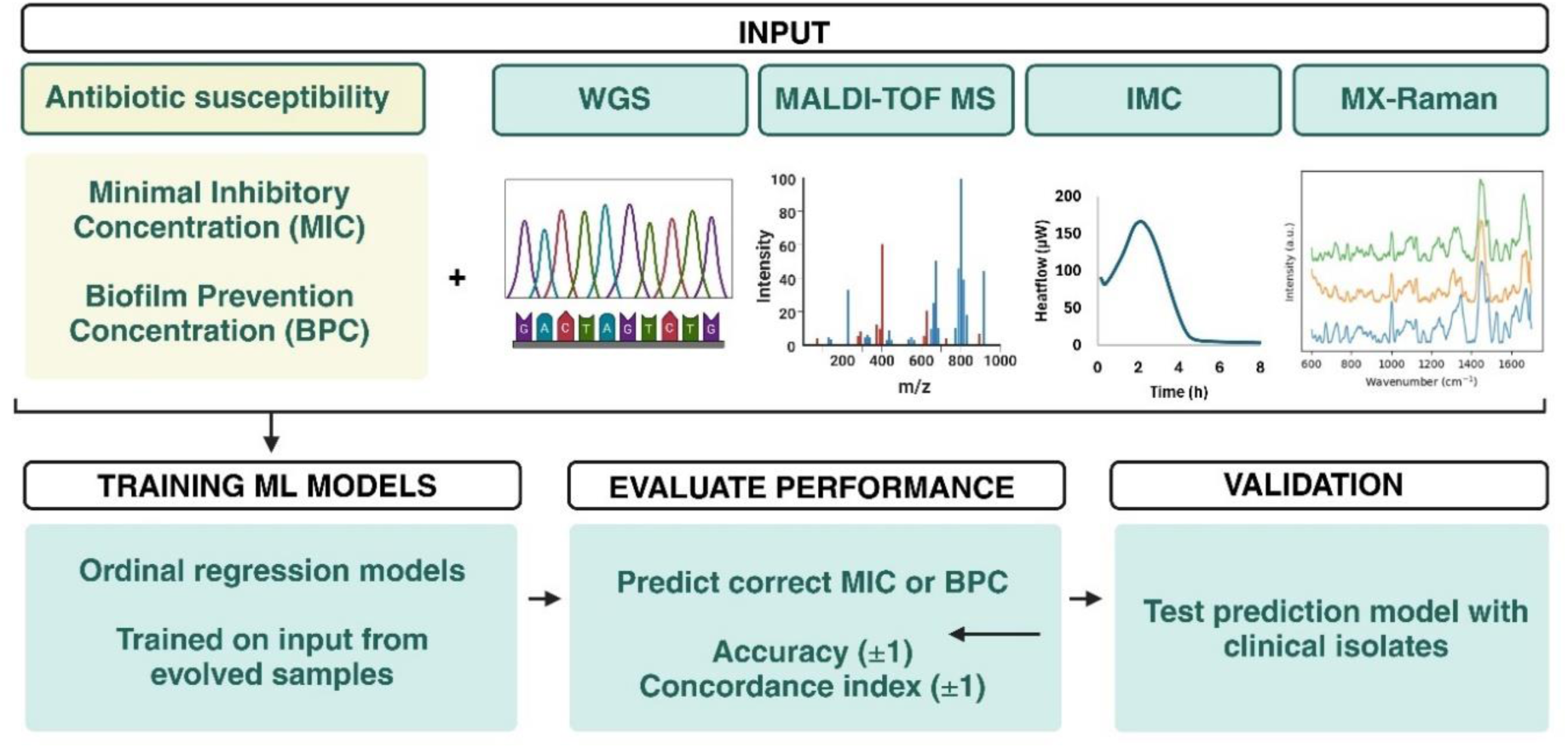
Overview of the workflow of the present study. The minimal inhibitory concentration (MIC) and the biofilm prevention concentration (BPC) of tobramycin were determined for 46 experimentally evolved *P. aeruginosa* strains (derived from six *P. aeruginosa* reference strains). These isolates were characterised using four analytical approaches, i.e. whole-genome sequencing (WGS), matrix-assisted laser desorption/ionization-time of flight mass spectrometry (MALDI-TOF MS), isothermal microcalorimetry (IMC) and multi-excitation Raman spectroscopy (MX-Raman). The resulting data were used to train ordinal regression machine learning models to predict MIC or BPC values. Model performance was assessed by evaluating the accuracy (±1) and the concordance index (±1). After developing the prediction model, it was validated with an independent dataset of clinical *P. aeruginosa* isolates. Created in BioRender.

## Results and Discussion

### Antibiotic susceptibility of experimentally evolved *P. aeruginosa* biofilms in SCFM2

We used an experimental evolution approach to generate a collection of evolved *P. aeruginosa* isolates derived from six reference strains. For each *P. aeruginosa* strain, eight independent cultures were set up in SCFM2 (except for CF1 with six cultures), resulting in 46 lineages that were maintained for 15 cycles. Half of the lineages were exposed to tobramycin and the other half served as untreated controls. While the amount of CFU/mL remained stable in control lineages, it significantly increased over time in all tobramycin-exposed lineages (except for AA2-1) (Fig. S1). At cycle 15, the experiment was concluded and the MIC and BPC of tobramycin were determined (Fig. 2). The strains exposed to tobramycin during evolution showed significantly higher MIC and BPC values compared to those evolved without antibiotic exposure, indicating that exposure during evolution led to a reduced susceptibility to tobramycin (Fig. S2).

**Figure 2.**
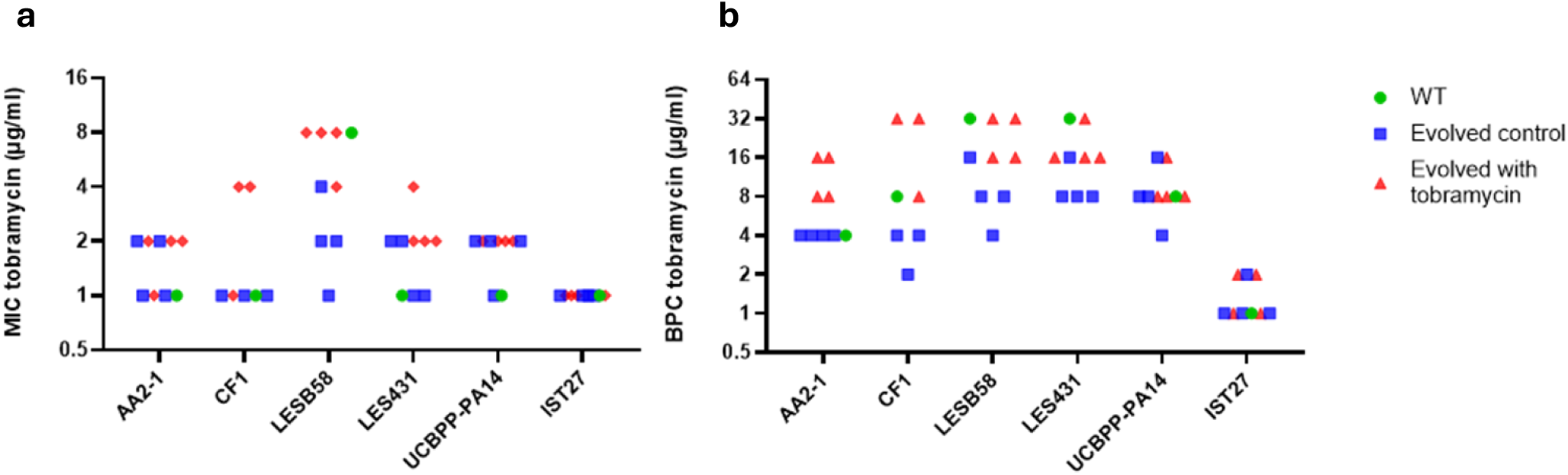
Antibiotic susceptibility of experimentally evolved *P. aeruginosa*. (**a**) Minimal Inhibitory Concentration (MIC) and (**b**) Biofilm Prevention Concentration (BPC) of tobramycin (µg/mL) for all evolved strains (blue squares: control, red triangles: exposed to tobramycin during evolution) and their respective wild-types (WT) (green dots). Data points represent the median value of three biological replicates per sample.

### Development of a machine learning model to predict antimicrobial susceptibility

#### Modelling and performance evaluation

A machine learning model was developed to predict the MIC or BPC of experimentally evolved strains based on WGS, MALDI-TOF MS, IMC or MX-Raman data. In what follows, a single sample denotes one evolved lineage, for which one prediction was made. Given the ordinal nature of the prediction targets (MIC or BPC values), an ordinal regression model was used. Prediction plots that visualise the probability at which each ordinal class was predicted, can be found in Fig. S3. Prediction quality was assessed in terms of accuracy and concordance index (C-index). The accuracy measures the percentage of samples for which the correct MIC or BPC was predicted. The C-index measures the probability that two randomly chosen predictions are ranked correctly. For both of these metrics, an additional version is computed that allows misprediction up to one 2-fold dilution step: accuracy^±1^ and C-index^±1^, respectively (for more information, see Methods).

To assess the prediction quality of the machine learning models, performances are compared to ‘random’ predictions. Here, ‘random’ performance consists of the score obtained when every prediction would be the MIC or BPC category that occurred the most often in the training dataset. More formally, the predictions *P*(*y* = *k* | *x*) correspond to the probability *P*(*y* = *k*), unconditioned on *x* . Hence, the score of a random prediction model corresponds to the setting where no relevant signal is present in the input data. This score was computed for both the accuracy and the accuracy^±1^. For the C-index and C-index^±1^, this random score is always 0.5, making these metrics more useful when comparing scores between MIC and BPC predictions. Models trained on MIC and BPC values have different ‘random’ performances as there are only four different MIC values present in our dataset (i.e. four ordinal categories), whereas the BPC dataset consists of six ordinal categories. A higher number of ordinal categories generally results in a more challenging prediction scenario and a lower random performance. Furthermore, the distribution of values also affects the random performance; evenly distributed categories make it harder to guess the correct value, whereas skewed distributions can increase the random performance score. It is therefore mandatory to compare a model’s performance to the random performance score and a model’s performance should be judged by how much it exceeds the performance of a random model.

Statistical tests typically require repeated measurements of performance obtained from independent test data sets. Given the nature of the evaluation metrics, delivering a single estimate of performance on the whole data set, in this setting, it is impossible to test for statistically significant differences between the different methods without violating test assumptions ^32^. Instead, in what follows, model performances for different data sources are compared using the aforementioned notion of random predictions. Informally, a (higher) performance level above random indicates the more obvious presence of signals indicative of antimicrobial resistance in the data.

#### Whole-genome sequencing

Whole-genome sequences were obtained for 46 experimentally evolved *P. aeruginosa* strains. By comparing these sequences to that of the unevolved WT, the mutational landscape could be mapped (Table S1). Next, a machine learning model was developed to predict the MIC and BPC values based on the detected DNA variants. This unbiased approach integrates all observed DNA variants, enabling the model to estimate which mutations affect MIC and BPC values, without requiring prior knowledge of determinants of susceptibility. While this approach is effective for experimentally evolved strains with a known WT ancestor, it is not directly applicable to clinical isolates as the choice of a ‘WT’ reference genome is inherently ambiguous (or even impossible) in that case.

The model predicted the MIC from WGS data with an accuracy^±1^ of 89.13 % (random performance: 82.61 %), while the BPC was predicted with an accuracy^±1^ of 76.09 % (random performance: 69.57 %). The C-index^±1^ was 71.77 % and 80.16 %, for MIC and BPC predictions, respectively (Fig. 3, Table S2), indicating that the model is slightly better in predicting the BPC than the MIC from WGS data.

**Figure 3.**
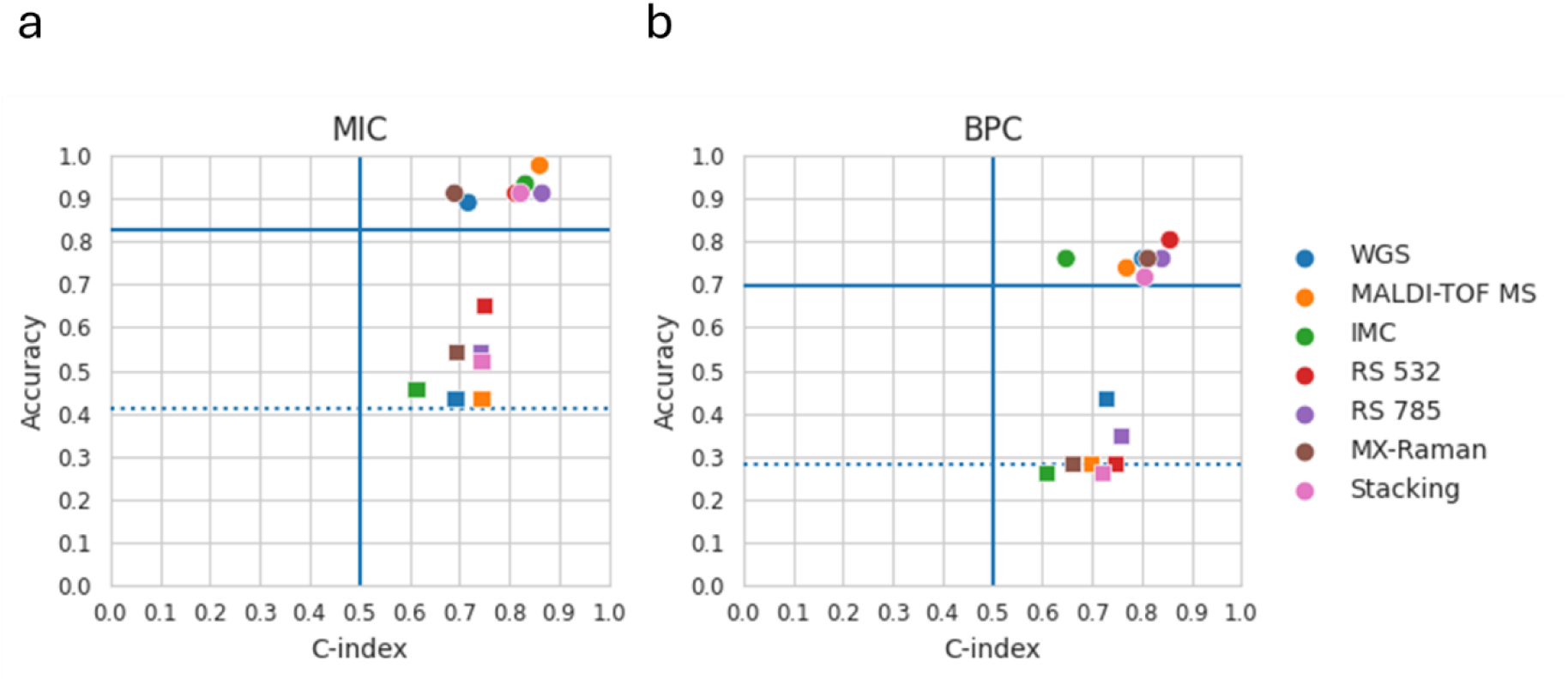
Performance of machine learning predictions based on various analytical approaches. (**a**) Scatterplot showing the performance for predicting the MIC (squares) or the MIC with an allowed margin of error of one 2-fold dilution (dots) (**b**) Scatterplot showing the performance for predicting the BPC (squares) or the BPC with an allowed margin of error of one 2-fold dilution (dots). The blue lines indicate scores based on random predictions. Horizontal blue dashes: random accuracy performance. Horizontal blue full line: random accuracy^±1^ performance. Vertical blue full line: random C-index and C-index^±1^.

WGS has been widely used to predict antimicrobial susceptibility, typically by screening for well-studied, known resistance genes ^18,33–36^. For instance, Eyre et al. used this approach to predict the MIC and MIC ±1 2-fold dilution of several antibiotics in *Neisseria gonorrhoeae* with accuracies of 53 % and 93 %, respectively ^33^. However, focusing only on known resistance genes poses a major limitation, as many tolerance and resistance mechanisms remain poorly understood. To address this challenge, we included all detected DNA variants as input for our machine learning model, regardless of prior knowledge on the role of specific gene products. Other studies have tackled these limitations by using entire genome sequences as input for prediction algorithms ^37–39^. For example, Nguyen et al. used this method to predict MIC values ±1 2-fold dilution for several antibiotics in *Salmonella*, achieving an accuracy^±1^ of 95 %, which is comparable to our results ^37^. In the present study, using entire genome sequences as input for the machine learning model was not possible due to the high-dimensional nature of this approach, which required a large sample size to be effective. Instead, we used DNA variants detected by mapping the entire genome sequences against a reference genome as input for the machine learning model. Nonetheless, integrating entire genome sequences with machine learning models would be an ideal strategy for future studies with larger datasets.

#### MALDI-TOF MS

MALDI-TOF mass spectra were collected for 46 experimentally evolved strains (Fig. 4a), and a machine learning model was developed to predict MIC or BPC values directly from these spectra, using them as unique spectral fingerprints. The model predicted the correct MIC with an accuracy^±1^ of 97.83 % (random performance: 82.61 %), while the BPC was predicted with an accuracy^±1^ of 73.91 % (random performance: 69.57 %). The model obtained a C-index^±1^ of 86.12 % and 76.92 % for MIC and BPC predictions, respectively (Fig. 3, Table S2). These results indicate that MALDI-TOF mass spectra allow a more precise prediction of the MIC than of the BPC. Overall, these results demonstrated that MALDI-TOF mass spectra can provide valuable insights into antibiotic susceptibility.

**Figure 4.**
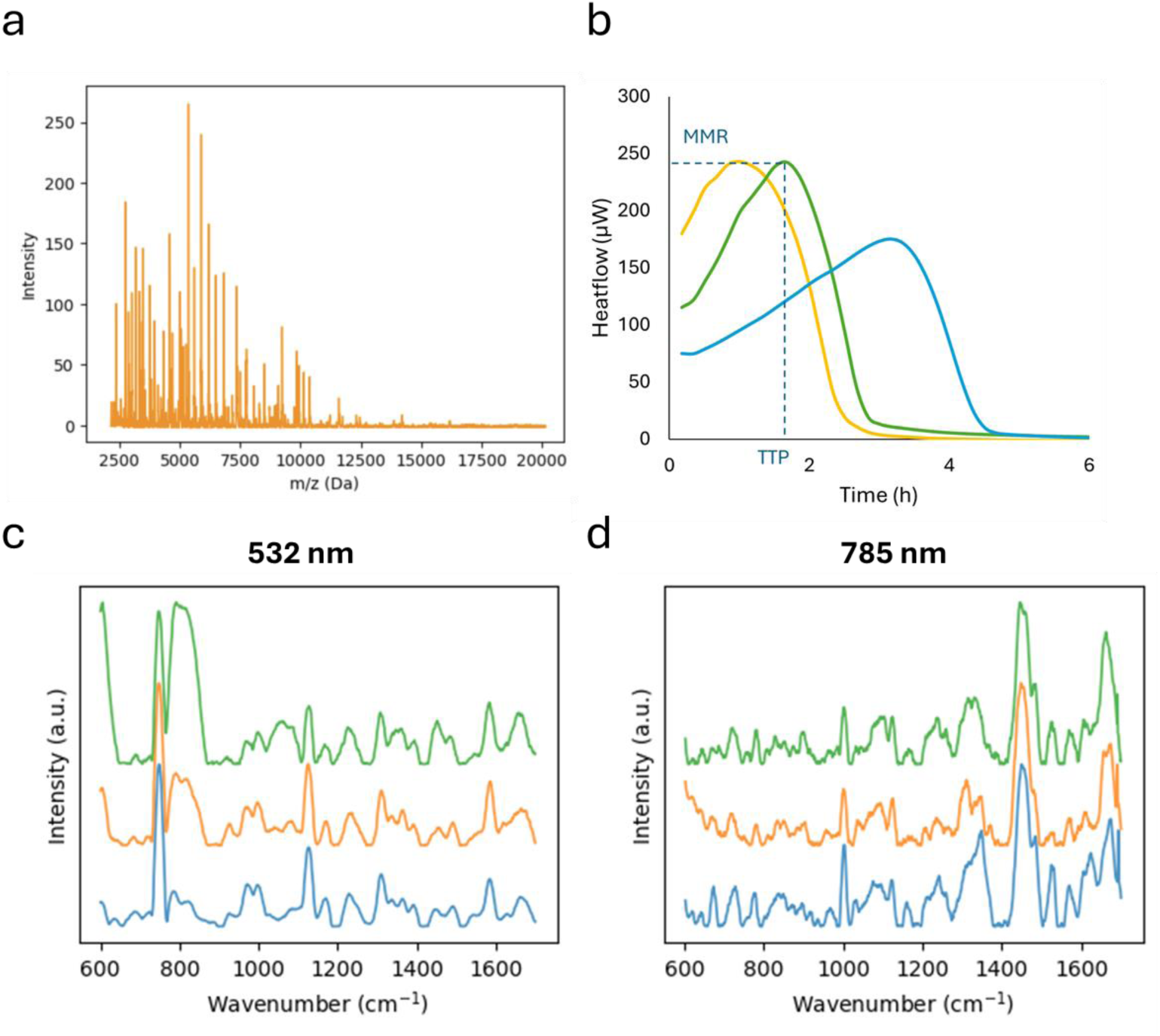
Examples of data input for the machine learning model. Data is shown of evolved *P. aeruginosa* AA2-1 L26 (orange) and L32 (green), and LES B58 L08 (blue). (**a**) MALDI-TOF mass spectra (**b**) Thermograms measuring the heatflow (µW) over time. Several parameters can be derived from thermograms, such as the time to peak (TTP) and the maximum metabolic rate (MMR) (**c**) Raman spectra obtained at excitation wavelength 532 nm (**d**) and 785 nm.

Recent studies have demonstrated the potential of machine learning models in classifying isolates as susceptible or resistant based on mass spectra ^23,40–42^. For example, Weis et al. classified *Staphylococcus aureus* isolates as susceptible or resistant with an area under the receiver operating characteristic curve (AUROC) of 80 % ^23^. Similarly, Nguyen et al. classified *P. aeruginosa* isolates resistant to ceftazidime/avibactam and ceftolozane/tazobactam, achieving AUROCs of 86.9 % and 85.6 %, respectively ^40^. The AUROC metric is equivalent to the C-index used in our study, but is specific to binary predictions. These studies predict whether the MIC is above or below a certain breakpoint (EUCAST or CLSI), but as such breakpoints are not yet established for biofilms, it is currently not possible to translate a BPC value into a classification as ‘resistant’ or ‘susceptible’. Our study addresses this limitation by training machine learning models to predict exact BPC values from the mass spectra, enabling compatibility with potential future biofilm breakpoints based on epidemiological cut-off values or other measures ^13^. Using this approach, our models obtained a C-index^±1^ of 86.12 % for MIC predictions, comparable to the aforementioned studies, and a C-index^±1^ of 76.92 % for BPC predictions.

Optimizing MALDI-TOF MS for AST could offer several advantages, as these devices are already widely used in hospitals for pathogen identification. Additionally, the technique has a high throughput, allowing the analysis of up to 96 samples in just minutes, making it a fast and efficient tool in clinical settings.

#### Isothermal microcalorimetry

In IMC, the heat produced by microbial metabolic processes is measured in real time. Using this method, we determined the metabolic profiles of 24-hour old biofilms in SCFM2 for all experimentally evolved strains (Fig. 4b). A machine learning model was trained to predict MIC and BPC values based on thermograms obtained with IMC. The model predicted the correct MIC with an accuracy^±1^ of 93.48 % (random performance: 82.61 %) and the BPC with an accuracy^±1^ of 76.09 % (random performance: 69.57 %). The model obtained a C-index^±1^ of 83.25 % and 64.78 %, for MIC and BPC predictions, respectively (Fig. 3, Table S2). These results suggested that IMC data were more predictive for the MIC than for the BPC. Nonetheless, both MIC and BPC predictions outperformed random predictions, indicating that even without antibiotic exposure, metabolic profiles of biofilms can reveal meaningful insights into their antibiotic susceptibility. Previous studies that explore the use of IMC for AST typically measured the metabolic activity of planktonic cultures or biofilms exposed to antibiotics ^25–27,43^. These approaches mirror the principle of broth microdilution MIC determination, but benefit from IMC’s high sensitivity (detection limit: 10^4^ CFU/mL ^44^) leading to faster results. Using this approach, Tellapragada et al. predicted the MIC of amikacin in planktonic *P. aeruginosa* with an essential agreement (number of results that were within one doubling dilution of the matching MIC determined by reference methods) of 97.4 % ^25^. In contrast, our setup measures untreated biofilms, eliminating the need to test multiple antibiotic concentrations and increasing throughput. To our knowledge, this is the first study that combines IMC data from untreated biofilms with machine learning algorithms to predict the MIC or BPC.

#### Raman spectroscopy

Raman spectroscopy provides a rapid and label-free method to obtain specific molecular fingerprints. In the present study, biofilms were grown overnight in SCFM2, and the resulting pellet from that overnight culture was applied to a fused quartz slide for analysis under a Raman microscope at two different excitation wavelengths, i.e. 532 and 785 nm (Fig. 4c, 4d). The spectral data from these individual wavelengths were also combined to create multi-excitation Raman spectra (MX-Raman) ^29^.

Using both single-wavelength (RS 532 nm or RS 785 nm) and MX-Raman spectra, a machine learning model was trained to predict MIC and BPC values. For MIC predictions, all three input types – RS 532 nm, RS 785 nm and MX-Raman – achieved an accuracy^+-1^ of 91.30 % (Fig. 3, Table S2). However, RS 785 nm yielded the highest C-index^±1^ (86.60 %), followed by RS 532 nm (81.34 %), and MX-Raman with the lowest C-index^±1^ (69.06 %). While Lister et al. previously reported that combining multiple wavelengths improved bacterial strain identification using a support vector machine (SVM) model, our results do not show a clear advantage for MX-Raman in susceptibility predictions ^29^.

For BPC predictions, values for accuracy^±1^ were similar across methods; i.e. 80.43 % for RS 532 nm and 76.09 % for RS 785 nm and MX-Raman. Likewise, C-index^±1^ scores were similar across methods (85.63 %, 84.01 % and 81.16 % for RS 532 nm, RS 785 nm and MX-Raman, respectively). Overall, the C-index^±1^ scores for both MIC and BPC predictions were quite comparable, except for MX-Raman’s notably lower performance on MIC predictions (69.06 % vs. 81.16 % for BPC). Nonetheless, this score is still substantially above random (50 %), suggesting that Raman spectroscopy is a good predictor for both MIC and BPC.

When the Raman spectra from samples sharing the same MIC or BPC value were averaged, distinct differences in peak intensities emerged among the resulting group averages, correlating with the respective MIC or BPC values (Fig. S4).

Previous studies have demonstrated the potential of Raman spectroscopy for differentiating resistant and susceptible bacteria based on unique spectral signatures ^30,31^. Lister et al. classified resistant bacterial strains with 100 % accuracy and susceptible strains with 98.89 % accuracy using an SVM model ^30^. Similarly, Lu et al. distinguished a susceptible *Acinetobacter baumannii* strain from five resistant strains with 99.92 % accuracy using a random forest model, though their approach could not identify to which antibiotics the strains were resistant ^31^. Our study is the first to combine Raman spectroscopy with machine learning to predict BPC values, demonstrating its potential as a valuable tool in biofilm susceptibility testing.

### Combining data from different analytical approaches does not increase the performance of MIC or BPC predictions

To evaluate if there is an added value in combining results from multiple analytical approaches, a stacking model that integrated data from all sources was used. This stacking approach used predictions based on the different types of data as inputs for a final model ^45,46^. Using the stacking model, MIC values were predicted with an accuracy^±1^ of 91.30 % (random performance: 82.61%), while the highest individual performance was achieved with MALDI-TOF (accuracy^±1^ of 97.83 %). A C-index^±1^ of 82.30 % was obtained with the stacking model, while the best-performing individual technique (RS 785 nm) had a C-index^±1^ of 86.60 %. For BPC predictions, the stacking model achieved an accuracy^±1^ of 71.74 % (random performance 69.57 %), whereas the highest accuracy^±1^ was obtained with RS 532 nm (80.43 %). The C-index^±1^ for BPC predictions with the stacking model was 80.57 %, while the best-performing method (RS 532 nm) achieved a C-index^±1^ of 85.63 % (Fig. 3, Table S2).

Overall, these results showed that combining data sources through stacking did not lead to a higher performance than the best-performing individual data source alone (Fig. 3). Nonetheless, the stacking model can provide insights into complementary data sources by analysing learned ordinal model coefficients β ∈ ℝ^*f*×1^ (Fig. S5). For both the MIC and BPC predictions, we observed highly positive coefficients for MALDI-TOF MS and WGS. The fact that both techniques showed a high variable importance indicated that they provided complementary, non-overlapping information. This finding suggested that, while we could not establish a performance gain with this combination in this study, future work with different datasets or additional refinement may reveal that combining WGS and MALDI-TOF MS data can further enhance prediction accuracy.

### Validation of machine learning model with clinical isolates

The prediction model was initially trained on data from experimentally evolved strains. To evaluate its ability to predict the MIC or BPC of clinical *P. aeruginosa* isolates, we analysed MALDI-TOF MS, IMC and MX-Raman data obtained from 30 CF isolates.

For MIC predictions, the best performance was observed for IMC, with an accuracy^±1^ of 80 % (random performance: 60 %) and a C-index^±1^ of 86.90 % (Fig. 5, Table S3). Predictions based on MALDI-TOF MS achieved an accuracy^±1^ of 76.67 % and a C-index^±1^ of 58.93 %. Using RS 532 nm, the accuracy^±1^ was 63.33 %, but the C-index^±1^ of 10.71 % was far below the random predictions of 50 %. RS 785 nm had an accuracy^±1^ of 60 % - equal to random performance – while its C-index^±1^ reached 53.57 %, only slightly exceeding the random performance of 50 %. MX-Raman achieved an accuracy^±1^ of 60 % - equal to random performance – and a C-index^±1^ of 47.02 %, which fell below random predictions.

**Figure 5.**
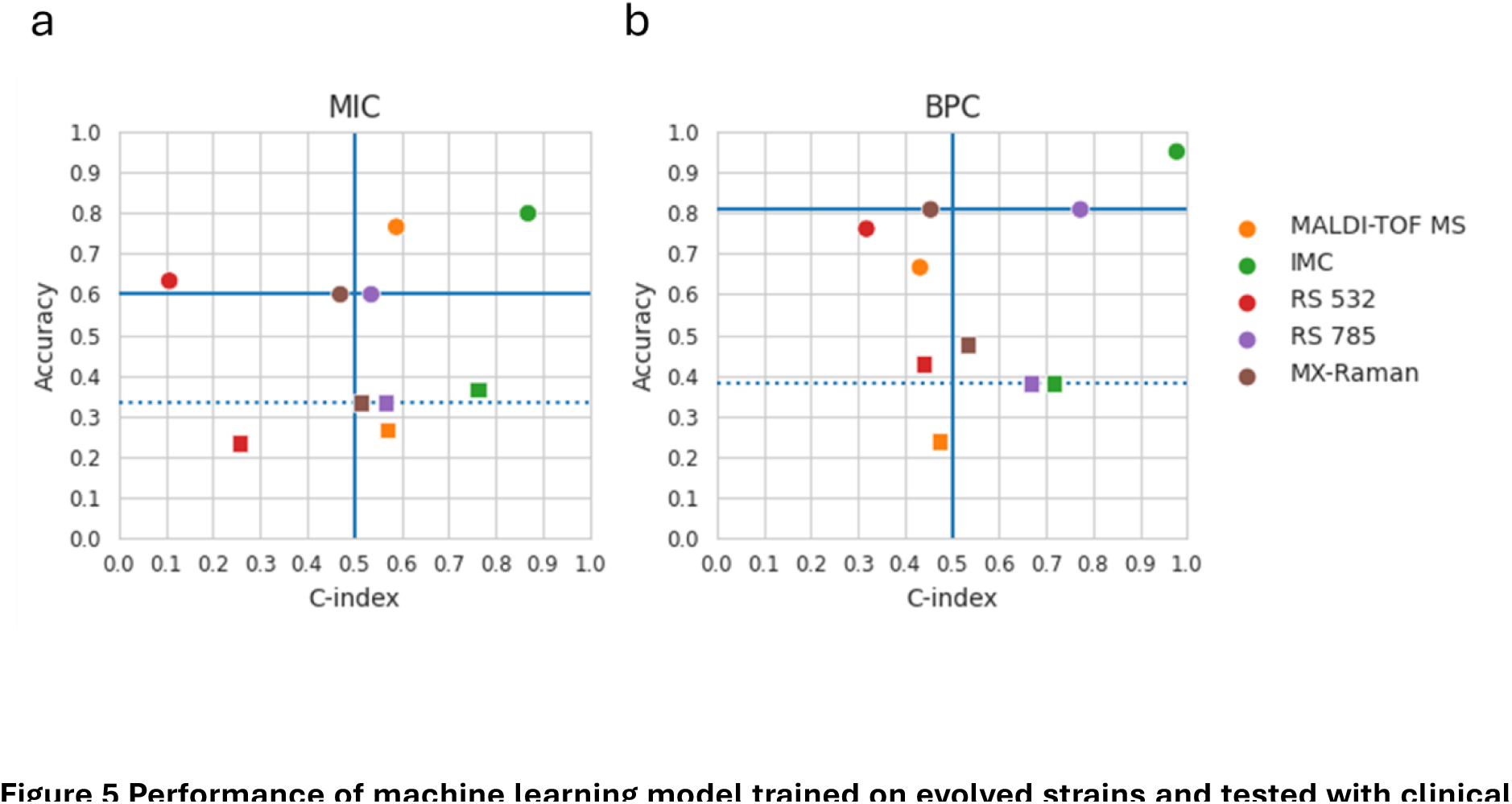
Performance of machine learning model trained on evolved strains and tested with clinical isolates. **(a)** Scatterplot showing the performance of predicting MIC (squares) or the MIC with an allowed margin of error of one 2-fold dilution (dots) **(a)** Scatterplot showing the performance of predicting BPC (squares) or the BPC with an allowed margin of error of one 2-fold dilution (dots). The blue lines indicate scores based on random predictions. Horizontal blue dashes: random accuracy performance. Horizontal blue full line: random accuracy^±1^ performance. Vertical blue full line: random C-index and C-index^±1^.

For BPC predictions, IMC again performed best, achieving a very high accuracy^±1^ of 95.24 % (random performance: 80.95 %) and C-index^±1^ of 97.73 %. Meanwhile, MALDI-TOF MS, RS 532 nm and MX-Raman all performed at or below random performance levels, with accuracy^±1^ scores of 66.67 %, 76.19 % and 80.95 %, respectively, and C-index^±1^ scores of 43.18 %, 31.82 % and 45.45 %. RS 785 nm achieved the second-highest C-index^±1^ of 77.27 % (well above the 50 % random prediction level), although its accuracy^±1^ of 80.95 % was the same as that of random predictions.

In summary, for MIC predictions, IMC and MALDI-TOF MS were the only techniques scoring above random predictions for both accuracy and C-index, with IMC showing the best performance. For BPC predictions, only IMC performed above random predictions for both accuracy and C-index. These results indicate that IMC data have the highest predictive power when it comes to predicting antimicrobial susceptibility, particularly for predicting the BPC. In contrast, the other techniques showed reduced performance when models trained on data from evolved strains were externally validated with clinical isolates. This discrepancy highlights the challenges of translating model performances across different datasets, a limitation also observed in other studies. For instance, Weis et al. and Ren et al. reported reduced performance when isolates from one dataset were tested on models that were trained on another dataset ^23,41^.

We hypothesize that the IMC data were easier for the model to interpret because the thermograms displayed clear trends (Fig. S6). As the MIC or BPC increased, the time to peak (TTP) also increased, while the maximum metabolic rate (MMR) generally decreased. This particularly resulted in a high C-index, as this is a ranking-based metric and the thermograms exhibited a clear ranking pattern. Despite visual differences between thermograms of clinical isolates and those of the experimentally evolved strains (Fig. S6), the model was still able to interpret them effectively. Furthermore, the observed IMC changes related to susceptibility are consistent with recent findings linking changes in bacterial metabolism to reduced susceptibility ^24,47^.

### Learned model coefficients suggest an important role for Anr and NppA1A2BCD mutations in tobramycin susceptibility

All experimentally evolved strains were genome-sequenced and compared to their ancestral wild-type (WT) strain. The observed mutations are listed in Table S1. Overall, we found that many global regulators and two-component systems (TCS), such as *lasR, pqsR, bfmR/bfmS* and *gacS/gacA* ^48,49^ acquired mutations. Additionally, numerous mutations were observed in proteins involved in 3’,5’-cyclic diguanylic acid (c-di-GMP) metabolism, including *wspF, dipA, morA* and *rbdA*, all of which play a role in biofilm formation ^50^. Interestingly, only one gene that has a known association with tobramycin resistance, i.e. the efflux pump regulator *mexZ* ^51^, was mutated in two tobramycin-evolved lineages.

Learned model coefficients can provide valuable insights into which genes might be correlated with increased MICs and BPCs. However, these correlations should be interpreted with caution, as some strongly correlated mutations may not directly correspond to causal biological mechanisms. For instance, two mutations might occur together within a single isolate with a high MIC. In such cases, it is possible that only one mutation is the actual causal (i.e. biologically relevant) factor driving the higher MIC value, while the other mutation merely co-evolved. Despite this, the model might estimate both mutations as being correlated with an increased MIC, even though only one exerted a true biological effect. Therefore, careful consideration is necessary when interpreting these results. Nevertheless, in large datasets, learned model coefficients can serve as valuable tools for identifying candidate genes that merit further investigation.

The model assigned a highly positive coefficient to mutations in *anr*, an anaerobic transcriptional regulator that controls the expression of genes essential for survival in low-oxygen environments. Previous studies have shown that regulation of the anaerobic respiratory pathway can reduce intracellular accumulation of aminoglycosides, contributing to adaptive resistance in *P. aeruginosa* ^52^. In the present study, mutations in *anr* were identified in CF1 populations evolved in the presence of tobramycin. Isolates with the amino acid substitution Phe→Ser 107 show a reduced susceptibility to the aminoglycosides tobramycin, gentamicin and amikacin. Compared to the CF1 WT (with an MIC of 1, 2 and 2 µg/mL for tobramycin, gentamicin and amikacin, respectively), the CF1 *anr* mutant displayed increased MICs of 4, 8 and 16 µg/mL for these antibiotics.

Several genes within the same cluster, encoding the ABC transporter NppA1A2BCD, also exhibited highly positive coefficients (Fig. S7). A previous study has shown that in *P. aeruginosa* NppA1A2BCD is required for the uptake of peptidyl nucleoside antibiotics ^53^. However, while we hypothesized that mutations in the Npp transporter also reduce uptake of the aminoglycoside tobramycin, we observed no differences in MIC or BPC of tobramycin between PA14 WT and the *ΔnppBCD* knockout mutant (MIC= 2 µg/mL, BPC= 8 µg/mL). When evaluating the minimal biofilm inhibitory concentration (MBIC) of tobramycin, a two-fold increase was observed in the knockout mutant (16 µg/mL in the PA14 WT vs. 32 µg/mL in the *ΔnppBCD* strain).

Other mutated genes with high positive coefficients include *rbdA* (PA0861), *bfmR* (PA4101) and *pmrA* (PA4776), although these were not experimentally investigated in the present study. The *rbdA* gene encodes a regulator of biofilm dispersal with phosphodiesterase (PDE) activity, and its inactivation in PAO1 has been demonstrated to lead to hyperbiofilm formation ^54^. The *bfmR* gene encodes a two-component system response regulator (TCS RR) that regulates genes involved in biofilm maturation and mediates the transition from acute to chronic virulence ^55^. Mutations in *bfmR* may indirectly influence antibiotic susceptibility by altering biofilm development or virulence pathways. The TCS RR *pmrA* regulates genes responsible for modifying the bacterial outer membrane (OM). Mutations in *pmrA* confer resistance to polymyxins by promoting OM modifications that reduce binding of cationic antibiotics ^56^. Furthermore, mutations in the PmrAB TCS have been shown to elicit cross-resistance to aminoglycosides ^57^.

## Conclusion

Using WGS, MALDI-TOF MS, IMC and RS data obtained from a collection of experimentally evolved *P. aeruginosa* strains, we were able to train machine learning algorithms that allowed to predict the MIC and BPC of tobramycin. All analytical approaches used demonstrated a predictive power that was higher than that of random predictions, confirming that each data type contained relevant information about antimicrobial susceptibility. We showed that an unbiased approach to predict susceptibility was possible with MALDI-TOF MS, IMC and MX-Raman data, as these methods do not require prior knowledge of mechanisms of susceptibility. Machine learning models successfully identified patterns in spectral data and thermograms, even without exposing the bacterial strains to antibiotics. This proof-of-concept study highlights the potential of alternative methods for predicting MIC and BPC values. Future research should focus on expanding datasets to include more strains and a wider range of MIC and BPC values across multiple antibiotics to further validate these promising results.

## Methods

### Bacterial strains, culture conditions and chemicals

The following *P. aeruginosa* strains were used: AA2-1 (a *lasR+* isolate derived from strain AA2; LMG 27630), LES B58 (LMG 27622), LES 431 (LMG 27624), UCBPP-PA14 (LMG 27639), IST27 (LMG 27643) ^58^ and CF1 (a Danish CF isolate belonging to sequence type 560). Strains derived from these wild type (WT) strains through experimental evolution are listed in Table S4. In addition, isolates recovered from chronically infected CF patients at Ghent University Hospital were used (Table S4); the collection of these strains was approved by the Ethics Committee of Ghent University Hospital, registration number B670201836204. The PA14 *ΔnppBCD* knockout mutant was previously described ^53^. The CF1 *anr* mutant (Phe→Ser 107) was isolated in the present study from an experimentally evolved CF1 strain (Table S4). Bacteria were stored at -80 °C in 8 % DMSO or using Microbank vials (Pro Lab Diagnostics, Canada) and were cultured at 37 °C on Tryptone Soy Agar (TSA) plates and in Tryptone Soy Broth (TSB) (Neogen, UK). Stock solutions of tobramycin (TCI Europe, Belgium), gentamicin (Sigma-Aldrich, USA) and amikacin (Sigma-Aldrich, USA) at 10 mg/mL were prepared in MilliQ water (filter sterilized, PES, 0.22 µm, VWR, Belgium). Synthetic cystic fibrosis medium (SCFM2) was prepared as described before ^14^, with the modification that mucin was sterilized by autoclaving instead of UV exposure.

### Experimental evolution

Six *P. aeruginosa* reference strains were experimentally evolved under biofilm conditions for 15 cycles, with or without exposure to tobramycin (Fig. 6). For each strain, four control lineages and four tobramycin-treated lineages were included (except for CF1, three lineages each), resulting in 46 independently evolved lineages. To this end, overnight cultures of *P. aeruginosa* were diluted in SCFM2 to approximately 5 x 10^7^ CFU/mL. 100 µL of the resulting suspension was added to a flatbottom 96-well plate (VWR, USA) and incubated for 24 h at 37 °C under aerobic conditions (without shaking). After 24 h, biofilms were treated with 100 µL of tobramycin solution or 100 µL of SCFM2 medium (for untreated controls). Tobramycin concentrations were selected based on preliminary experiments that resulted in a 2-3 log reduction in the number of CFU in a 24 hour old biofilm; i.e. 16 µg/mL for CF1, 32 µg/mL for UCBPP-PA14 and IST27, 64 µg/mL for AA2-1 and LES 431 and 128 µg/mL for LES B58. After an additional 24 h incubation at 37 °C, biofilm aggregates were disrupted by vortexing (5 min, 900 rpm) (Titramax 1000, Heidolph Scientific Products GmbH, Germany) and sonication (40 kHz, 5 min) (Branson 3510, Branson Ultrasonics, USA) and the number of CFU/mL was quantified by plating on TSA. A frozen stock of each lineage was stored at -80 °C in cryovials with 8 % DMSO in TSB. An overnight culture was prepared by inoculating 5 mL TSB and incubating for 24 h at 37 °C while shaking. The next day, a new cycle was initiated by inoculating with 5 x 10^7^ CFU/mL from that overnight culture. This process was repeated for 15 cycles.

**Figure 6.**
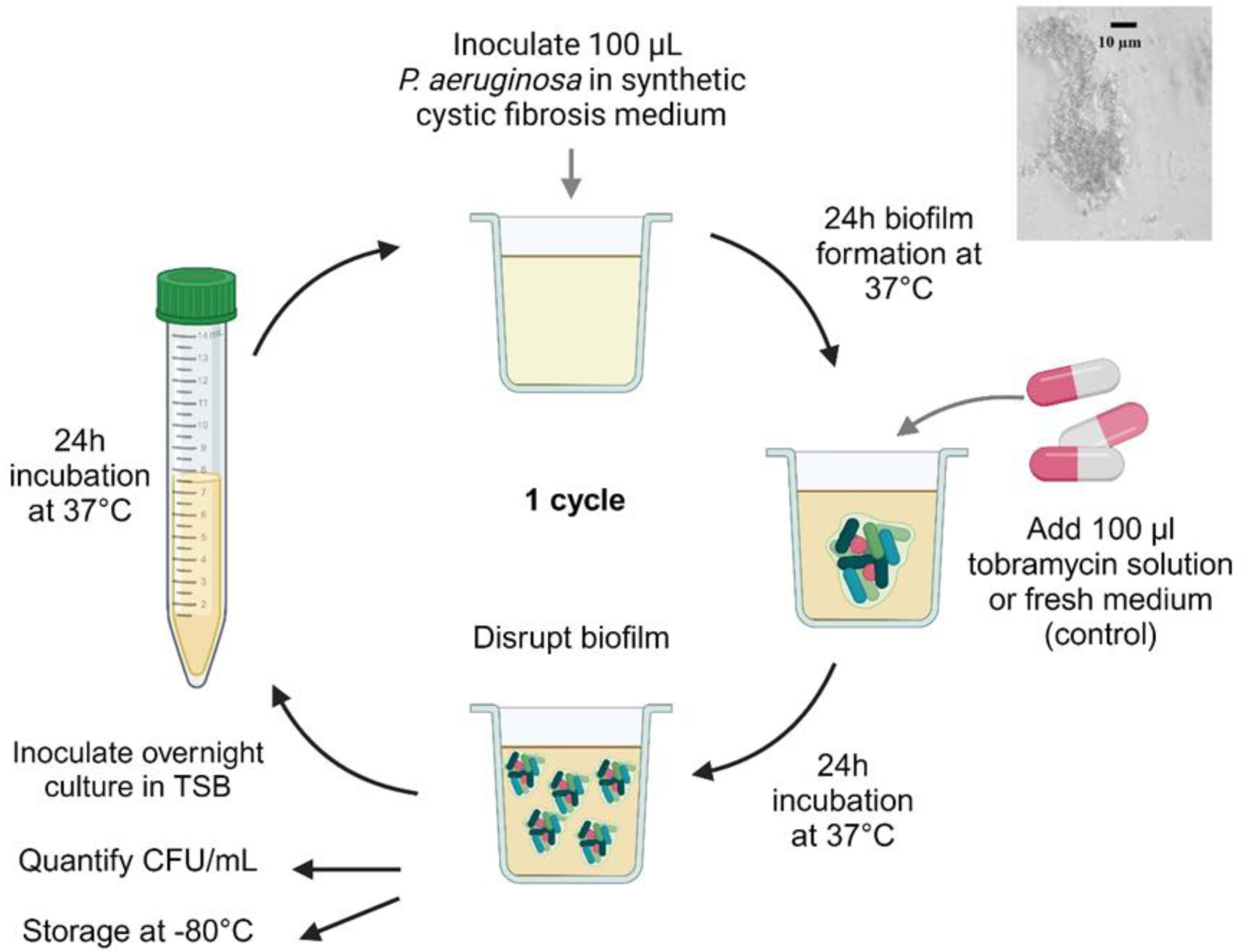
Experimental set-up of the biofilm evolution model in SCFM2. 100 µL of a *P. aeruginosa* culture is added to the well of a 96-well plate. After 24 h, suspended biofilm aggregates are formed and these are either treated with 100 µL of tobramycin, or 100 µL fresh SCFM2 medium is added. After 24 h, the biofilm is disrupted and a part of the bacterial suspension is used to inoculate an overnight culture to allow the start of a new cycle. The number of surviving cells after each cycle is quantified by plating on TSA. The samples are stored at -80 °C. Created with BioRender.

### Antibiotic susceptibility testing

MICs of tobramycin, gentamicin and amikacin were determined following the EUCAST guidelines using the broth microdilution method ^3^. MICs were defined as the lowest concentration of an antimicrobial agent that inhibited at least 90 % of microbial growth after 24 h of incubation. The BPC of tobramycin was determined using serial dilutions of the antibiotic in SCFM2 medium and inoculating bacteria at a final concentration of 5 x 10^7^ CFU/mL in SCFM2 ^10^. After 24 h incubation at 37 °C, the contents of the wells were plated and after 24 h of incubation colonies were counted. The BPC was defined as the lowest concentration of antimicrobial agent that prevented at least 90 % of biofilm growth compared to the growth control after 24 h of incubation. For selected isolates, the minimal biofilm inhibitory concentration (MBIC) of tobramycin was determined. The MBIC is the lowest concentration of tobramycin that resulted in at least 90 % reduction of biofilm growth compared to the growth control). To this end, 50 µL of a 5 x 10^7^ CFU/mL bacterial suspension in SCFM2 was added to round bottom 96-well plates. After 24 h incubation at 37 °C, double concentrated tobramycin solutions prepared in SCFM2 were added to each well. After 24 h of treatment at 37 °C, biofilms were disrupted by vortexing and sonicating (40 kHz, Branson 3510, Branson Ultrasonics, USA) the plate for 5 min each. The contents of the wells were plated on TSA, and after 24 h incubation at 37 °C, colonies were counted. All experiments were performed in biological triplicate.

### Statistical analysis

Statistical analysis was performed using IBM SPSS Statistics (version 29). The normality of the data was verified with a Shapiro-Wilk test. If the data were normally distributed, a two-sided independent samples *t*-test was performed to compare differences between two timepoints or two treatment groups. If the data were not normally distributed, a two-sided nonparametric Mann-Whitney U test was performed.

### DNA extraction

Overnight liquid cultures were centrifuged to obtain a pellet. The pellet was resuspended in 200 µL 10 mM TE-buffer (10 mM Tris-HCl pH 8, 1 mM EDTA pH 8), after which 100 µL was transferred to a 2 mL microcentrifuge tube containing approximately 500 µL acid-washed glass beads (≤106 μm; Sigma Aldrich, USA) and 500 µL lysis buffer (50 mM Tris-HCl pH 8, 70 mM EDTA pH 8, 1 % SDS) with 0.5 mg/mL Pronase (Roche, Germany). Samples were vortexed vigorously for 5-10 s, incubated at 37 °C for 30-60 min, and centrifuged at 13,000 rpm for a short spin. Following the addition of 200 µL of saturated ammonium acetate, samples were vortexed vigorously for 5-10 s and centrifuged again at 13,000 rpm for 2 min. To separate phases, 600 µL chloroform was added, and samples were vortexed horizontally for 5-10 s, and centrifuged for 5 min at 13,000 rpm. Then, 400 µL of clear aqueous top phase was transferred to an Eppendorf LoBind microcentrifuge tube (Eppendorf AG, Germany) containing 1 mL 100 % ethanol. The tubes were mixed by inversion to precipitate the DNA, followed by centrifugation at 13,000 rpm for 5 min. Afterwards, the supernatant was discarded and the pellet was washed with 500 µL 70 % ethanol. After a short spin, the ethanol was removed by pipetting and the pellet was air-dried. The dry pellet was dissolved in 300 µL low EDTA TE-buffer (10 mM Tris-HCl pH 8, 0.1 mM EDTA, 0.5 µg/mL RNase) and incubated at 37 °C for 60 min. DNA concentrations were determined using the BioDrop µLITE (BioDrop, UK).

### Whole-genome sequencing and data analysis

A PCR-free library preparation was performed using the NEBNext Ultra II Library Prep Kit for Illumina, following a size selection protocol using AMPure XP beads after adapter ligation. Samples were sequenced on the Illumina NextSeq 500 System, generating 75 bp single-end reads. The reads were analysed with CLC Genomics Workbench and mapped to reference genomes of *P. aeruginosa* AA2 (NZ_CP051547.1), LES B58 (NC_011770.1), LES 431 (NC_023066.1), UCBPP-PA14 (NZ_CP034244.1) or to the 28 contigs of IST27 (whole genome shotgun sequencing project MCMX01000001 to MCMX01000028), or the 36 contigs of CF1. Reads were mapped using a local alignment and filtered based on a 50 % length fraction and 80% similarity fraction. The basic variant detection tool was used to detect single nucleotide polymorphisms (SNPs) with a minimum frequency of 10 %, minimum count of 5, minimum quality of 20 and minimum forward/reverse balance of 0.3. All SNPs were manually screened to remove false positives. Insertions and deletions were detected using the InDels and Structural Variants tool, with a minimum sequence complexity of 0.2 and a minimum count of 5. Entries meeting minimum requirements were further manually filtered for false positives derived from sequencing and mapping errors. The raw reads generated in this study are available in the ArrayExpress database under the accession number E-MTAB-11894.

### MALDI-TOF mass spectrometry

Pure cultures were plated in biological duplicate on TSA. Confluent growth was collected with a sterile loop and suspended in 300 µL MQ. After vortexing, 900 µL of 100 % ethanol was added and the tubes were homogenized by inversion. After centrifuging for 3 min at 4 °C, the ethanol was discarded and evaporated. Next, the cell pellet was suspended in 40 µL formic acid and the samples were vortexed. Then, 40 µL acetonitrile was added and the samples were vortexed. After centrifugation, 1 µL supernatant (containing the protein extract) was spotted in duplicate on the target plate. Subsequently, 1 µL matrix solution (10 mg/mL alpha-cyano-4-hydroxycinnamic acid) was spotted on the plate. A bacterial test standard (BTS) was included for calibration. Mass spectra were obtained using the Biotyper Microflex LT/SH MALDI-TOF MS system (Bruker Daltonik GmbH, Germany). The experiments were performed in biological and technical duplicate.

### Isothermal microcalorimetry

Overnight cultures were diluted in SCFM2 medium to a final concentration of 5 x 10^7^ CFU/mL, and 100 µL of the bacterial suspension was transferred to plastic inserts (calVials, Symcel, Sweden) for biofilm growth over 24 h at 37 °C. The next day, 100 µL of fresh SCFM2 medium was added to the biofilms. The plastic inserts were then transferred to titanium cups and microcalorimetric measurements were conducted using the calScreener device (Symcel, Sweden). The resulting thermograms (in which heat production is plotted over time) were analysed with calView 2.0 software (Symcel). All experiments were performed in technical duplicates.

### Raman spectroscopy

Bacterial cultures were grown overnight at 37 °C in SCFM2 medium and were centrifuged for 10 min at 4,000 rpm. Pellets were washed with ddH_2_O, and after another round of centrifugation the resulting pellet was applied to a fused quartz microscopic slide (UQG Optics, UK) and dried on a heater. Samples were excited with a 532 nm or a 785 nm lasers using the Renishaw InVia Raman Microscope (Renishaw, UK). Spectra were acquired over three accumulations with a 5 s exposure time. Cosmic rays were removed from all spectra using the Renishaw WiRe 5.5 software. Multi-excitation spectra were obtained by manually merging the 532 nm spectrum to the end of the 785 nm spectrum.

### Data preprocessing for machine learning

Four different data types were collected, i.e. DNA variants, MALDI-TOF mass spectra, thermograms, and Raman spectra. To prepare these data for machine learning modeling, data were preprocessed to fixed-length feature vectors. In what follows, a ‘sample’ denotes a single evolved lineage. For all data sources – aside from DNA variants – multiple technical replicates per sample were generated. In data preprocessing, technical replicates were considered independently, every technical replicate becoming a separate row in the final machine learning input matrix. All preprocessed data are available at the following GitHub repository: https://github.com/gdewael/biofilm-amr.

Considering all possible variants in the genome constitutes an infeasibly big feature space, and for that reason DNA variant data were collated on a per-gene basis. The DNA variant input for the machine learning model, hence, was defined as ‘whether a variant was found in gene X’, for any gene X. The feature space was further reduced by eliminating loci for which no variant was found in any sample. Further, as machine learning cannot learn generalizing patterns for features present in a single sample, genes with variants in a single sample were similarly eliminated. The resulting feature vector for every sample was of binary nature, with every feature consisting of a single gene, indicating whether a variant in it was found in said sample. For MALDI-TOF data, spectra were preprocessed according to standard practices ^23^: (1) square-root transformation of intensities, (2) smoothing with a Savitzky-Golay filter using a half-window size of 10, (3) baseline correction with the 20 iterations of the SNIP algorithm, and (4) trimming to the 2,000-20,000 Da range. To reduce the feature space, peaks were detected using the persistence transformation algorithm, keeping only the top 128 for every spectrum ^59^. A fixed-length feature vector was then obtained by placing the detected peaks in 3Da-interval bin features (ranging from 2,000-20,000Da, 6,000 bins overall). To retain congruence with DNA variant data, bins were binarized, hence, each feature in the input vector indicates whether a peak was found in a bin. Features were similarly eliminated if they were found in either no spectra, or only one spectrum in the entire dataset.

Raman spectra were preprocessed similarly to the MALDI-TOF mass spectra. Briefly, the following steps were performed: (1) square-root transformation of intensities, (2) smoothing with a Savitzky-Golay filter using a half-window size of 10, (3) baseline correction with the 20 iterations of the SNIP algorithm, and (4) trimming to the 600-1700 cm^-1^ range. After, peaks were similarly detected using the persistence transformation algorithm, keeping only the top 128 peaks for every spectrum. Peaks were then binned in 1 cm^-1^ intervals and binarized. Features were eliminated if they were found in either no, or only one spectrum in the entire dataset. For MX-Raman, the final feature representation constitute the features for the 532 nm and 785 nm spectra, taken together by concatenation.

Microcalorimetry data are presented as thermograms, i.e. heatflow (µW) over time. Heatflow measurements were taken at regular intervals in time (every 10 min), so these data are already in fixed-length feature format (every feature being the heatflow in µW at a specific time point). Data were normalized to a range between zero and one by dividing all heatflow numbers by 250 (the highest heatflow number encountered rounded up to ten).

### Machine learning modeling

The prediction targets (MIC or BPC values) are of ordinal nature: 1, 2, 4, … (µg/mL). For this reason, an ordinal regression model is used to predict MIC or BPC values based on one of the data types described above. More formally, let us denote an input data sample as *x* ∈ ℝ^*f*^, with *f* number of input predictors. Its corresponding MIC or BPC value is given by *y* ∈ {2^*n*^ | *n* ∈ ℤ}. The cumulative logistic link function ^60^, then, models *y* as a function of *x* as follows:

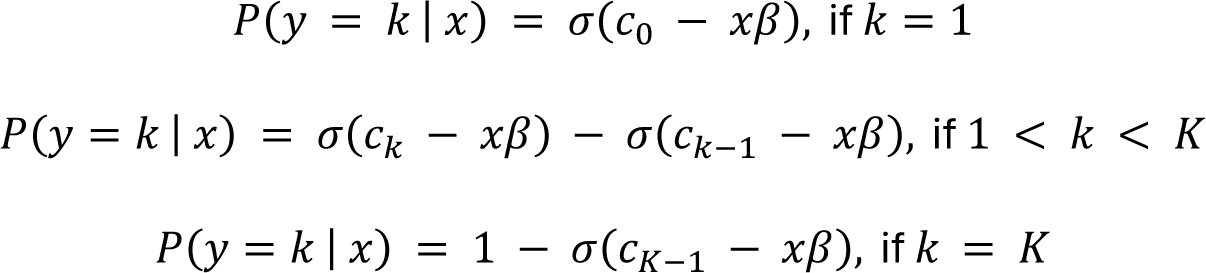

Where *k* denotes the ordinal classes numbered 1 to *k*. These ordinal classes map to the distinct MIC or BPC values in the data. For MIC, these correspond to: {1, 2, 4, 8}. For BPC values, these correspond to: {1, 2, 4, 8, 16, 32}. In essence, this model linearly transforms the input data to one dimension via learnable coefficients β ∈ ℝ^*f*×1^. This one-dimensional space is then ‘cut up’ into ordinal classes via learned (strictly increasing) cutpoints *c*_{1,…,*k*−1}_. The ordinal model can be used to either (1) predict the probability that a sample will have a certain MIC or BPC: *P*(*y* = *k* | *x*), or (2) predict a single MIC or BPC value for a sample: arg m ax_k_ *P*(*y* = *k* | *x*). Both the coefficients β and cutpoints *c*_{1,…,*k*−1}_ were jointly optimized to minimize the negative log-likelihood:

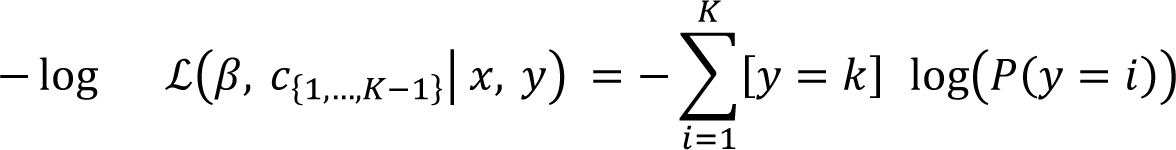

Models were trained using gradient descent. Every training iteration considered the full training data set as batch. All code to train models and fully reproduce all experiments is available at https://github.com/gdewael/biofilm-amr.

### Hyperparameter tuning

Models were tuned separately for every data source and target type (MIC or BPC). Hyperparameter tuning was performed through nested leave-one-out cross validation (CV). In case of technical replicates (present for all data sources, except DNA variant data), this data splitting scheme was adjusted to leave-one-group-out CV, with a group constituting all replicates corresponding to one sample. The outer CV loop served to obtain predictions. Hyperparameter tuning is performed for every iteration in the outer loop through an additional CV inner loop. Optimal hyperparameters are found through grid search, using the following possible values for hyperparameters: learning rate: {0.1, 0.5, 1}, number of iterations: {250, 750}, L2 regularization of coefficients: {0.0001, 0.001}. Quality of every hyperparameter configuration was determined using the concordance^±1^ index on every held-out sample of the inner CV. After every inner CV loop, a final model was re-trained using the optimal hyperparameters on the full validation and training set. This final model was used to make predictions for the sample left out of the outer loop.

For the Raman spectroscopy dataset, an exception was made to this procedure because of the size of the dataset (150 replicates per sample). Whereas other data sources used leave-one-(group)-out CV both in the inner and outer CV loops, for Raman spectroscopy, the inner (tuning) CV loop used 5-fold CV.

### Model evaluation

Model prediction quality was evaluated using the outer loop of a nested leave-one-out CV. To evaluate predictions for data sources with differing number of technical replicates on equal footing, predictions for technical replicates were collated to a single sample-level prediction. To obtain predicted probabilities at the sample level, a prediction was made for every technical replicate, and their probabilities for each ordinal category were averaged across replicates. All model evaluations occurred on these sample-level predictions.

Model prediction quality was assessed through the accuracy score, the accuracy^±1^ score, the concordance index, and the concordance^±1^ index. The accuracy score indicates how often the MIC/BPC category with the highest predicted probability was the correct one. The accuracy^±1^ score indicates how often a predicted MIC/BPC belonged to the correct ordinal category or one category higher or lower (i.e. the correct MIC/BPC or within one 2-fold dilution step, which is the accepted variability of phenotypic AST) ^2,37,61^. The concordance index evaluates the overall ranking quality of predictions. It evaluates every pair of samples with different MIC/BPC values and counts the proportion of pairs for which the higher MIC/BPC value also had a higher predicted MIC/BPC value. Because of its pairwise evaluation, it is numerically more stable in small sample sizes than the accuracy score. The concordance^±1^ extends this metric by only evaluating pairs of samples with MIC/BPC values differing by at least two 2-fold dilution steps. Here, the variant of the concordance index was applied that does not count ties as ‘half-correct’ ^62^. To test how much information the machine learning models learned from data, performances are compared to ‘random’ predictions. Here, ‘random’ performance consists of the score obtained when every prediction would be the MIC or BPC value that occurred most often in the training data set. This score was computed for both the accuracy and accuracy^±1^. For the concordance and concordance^±1^ indices, this score always corresponds to 0.5.

### External validation of models with clinical isolates

To externally validate the trained models, data obtained with 30 CF-derived *P. aeruginosa* isolates were used. The same training and tuning procedures as described above were used, the only difference being that in this case an external test set was used. Because of this, it was not necessary to perform nested cross validation to obtain unbiased estimates of model performance. Instead, only the previously described inner cross-validation loop was used to obtain optimal hyperparameters.

To establish random predictions for the clinical isolate test set, the same procedure as previously described was used, i.e. ‘random’ predictions are taken as the most frequently occurring class in the training data set (consisting of evolved strains).

### Combining data sources through stacking

To combine information from various data sources, stacking models were used. In stacking models, predictions from previous models are used as input for a second model. Per sample, to construct a single feature for every data source, the weighted sums of predicted probabilities of their previously trained models were computed: *x_j_* = ∑^*K*^_*i*=1_ *i* ⋅ *P_j_*(*y* = *i*), where *j* denotes one of the four used data sources. After data set construction, all stacking models were trained and tuned identically as previously described (i.e. using nested leave-one-(group)-out CV). As all previously trained models used leave-one-(group)-out CV as their splitting strategy, no specialized strategies were necessary to prevent leakage of information from training to evaluation sets.

### Data availability

All data necessary for supporting the findings of this study are enclosed in this manuscript. The raw sequencing reads generated in this study are available in the ArrayExpress database under the accession number E-MTAB-11894. The reference genomes used during sequencing analysis can be found in the GenBank database with the accession numbers NZ_CP051547.1 (AA2), NC_011770.1 (LES B58), NC_023066.1 (LES 431), NZ_CP034244.1 (UCBPP-PA14) and MCMX01000001 (IST27). All preprocessed data are available at the following GitHub repository: https://github.com/gdewael/biofilm-amr.

### Code availability

Code used in this study is available at https://github.com/gdewael/biofilm-amr.

## Acknowledgements

Part of this work was funded by the Ghent University Special Research Fund (grant BOF20/GOA/002). W.W. received funding from the Flemish Government under the “Flanders AI research program”. We thank dr. Daniel Pletzer (University of Otago, New Zealand) for providing the PA14 Δ*nppBCD* knockout mutant.

## Author contributions

T.C. developed the conceptual framework, supervised the project, and contributed to the manuscript. F.V. conducted the experiments, analysed the data, and wrote the manuscript with input from all authors. W.W. and G.D.W. managed the machine learning aspects of the study, with G.D.W. also contributing to the manuscript. F.V.N. made the whole-genome sequencing analysis possible. A.S. analysed the sequencing data and helped with the experimental evolution study. T.B. and M.L. provided expertise and assistance with the isothermal microcalorimetry analysis. J.S.W., S.M., C.H., N.H. and Y.C. contributed to the Raman spectroscopy analysis, while P.V. and M.C. assisted in the MALDI-TOF measurements.

