## Supplementary material for "Harnessing machine learning to predict antibiotic susceptibility in *Pseudomonas aeruginosa* biofilms": Supplemntal figures

### Supplementary figures

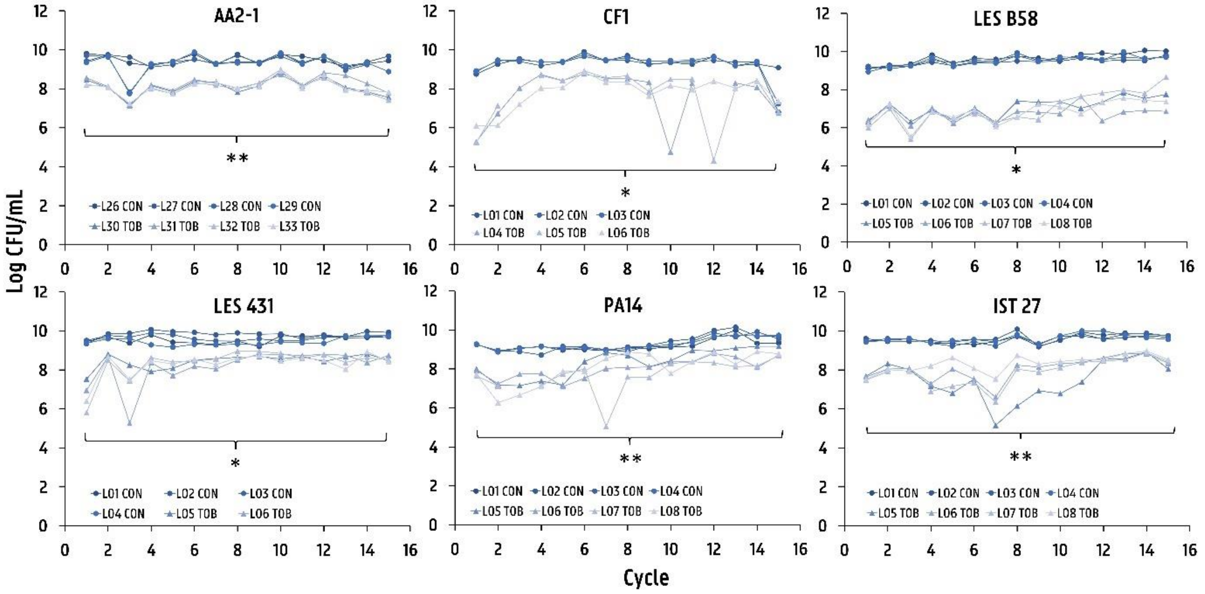


**Figure S1 Number of CFU/mL recovered after each cycle during experimental evolution.** Per strain, 4 independent lineages were included that were treated with tobramycin, and 4 lineages were left untreated, except for CF1, where 3 lineages each were included (per group, n=4, except for CF1 where n=3) (* *p*<0.05, ** *p*<0.01).

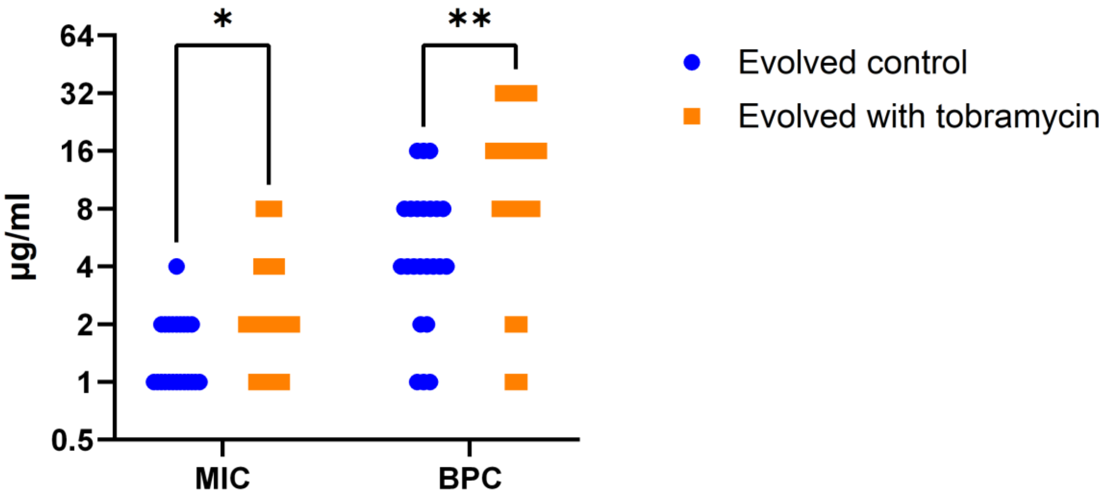


**Figure S2 Comparison of MIC and BPC values for tobramycin (µg/mL) between strains that evolved in the absence and in the presence of tobramycin** (per group, n=23) (* *p*<0.05, ** *p*<0.01)

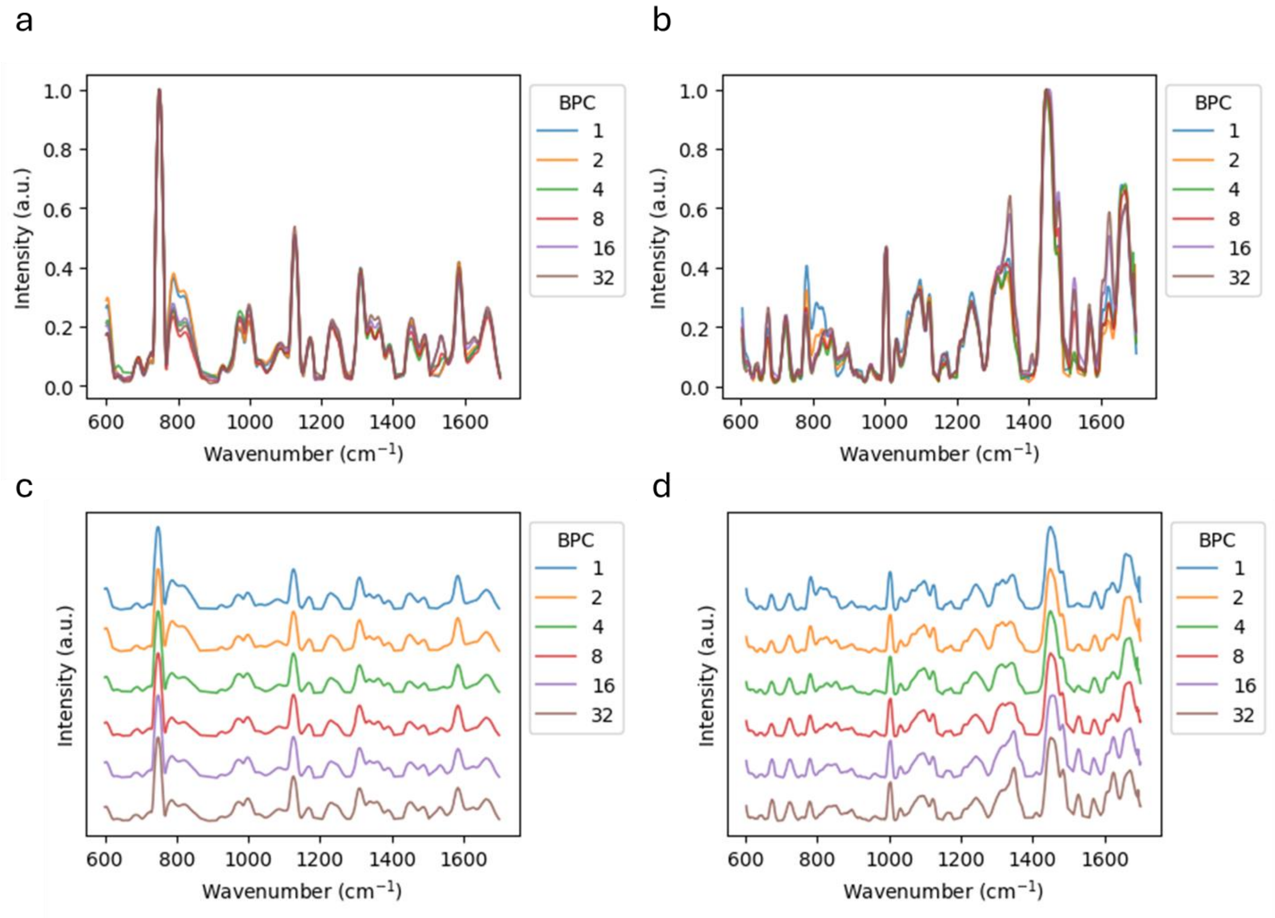


**Figure S4 Average Raman spectra of experimentally evolved strains grouped by BPC value.** Panels (a) and (b) show spectra acquired at 532 nm and 785 nm, respectively. Panels (c) and (d) display the same spectra offset for clarity, again at 532 nm and 785 nm, respectively. All spectra underwent the preprocessing steps described in "Data preprocessing for machine learning", except for peak detection.

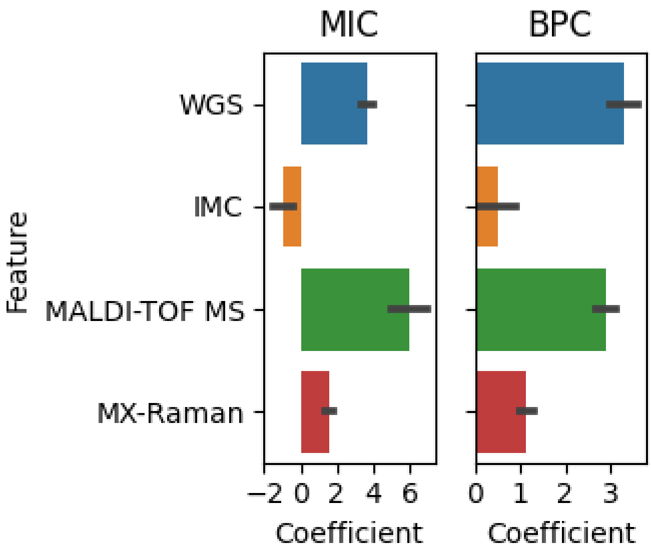


**Figure S5 Variable importances in the stacking model.** Coefficients refer to the linear transformation coefficients $\beta$ in the ordinal regression model. Error bars indicate standard deviation across 46 cross-validation folds. The left-hand and right-hand sides show coefficients for the model trained to predict MIC and BPC, respectively.

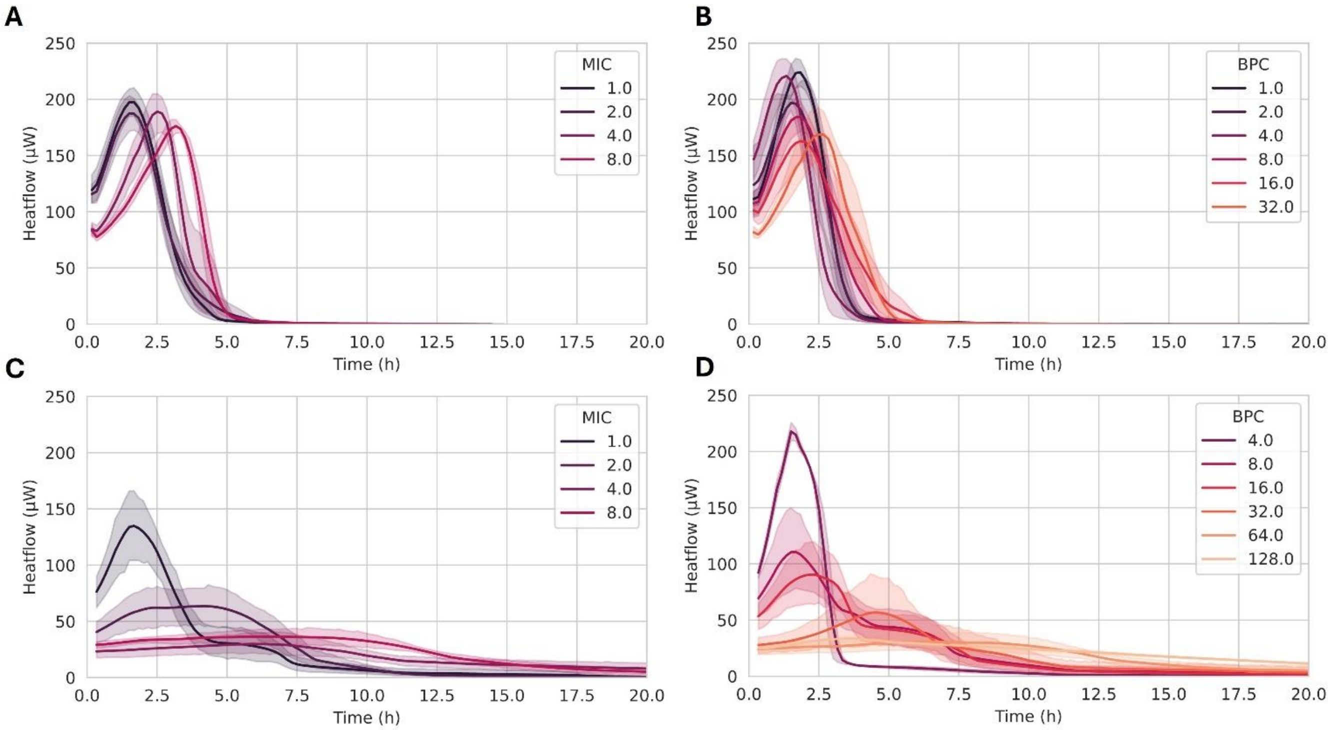


**Figure S6 Average thermograms grouped by MIC or BPC values.** Average thermograms for experimentally evolved strains (a) grouped by identical MIC values or (b) by identical BPC values. Average thermograms for clinical isolates (c) grouped by identical MIC values or (d) by identical BPC values.

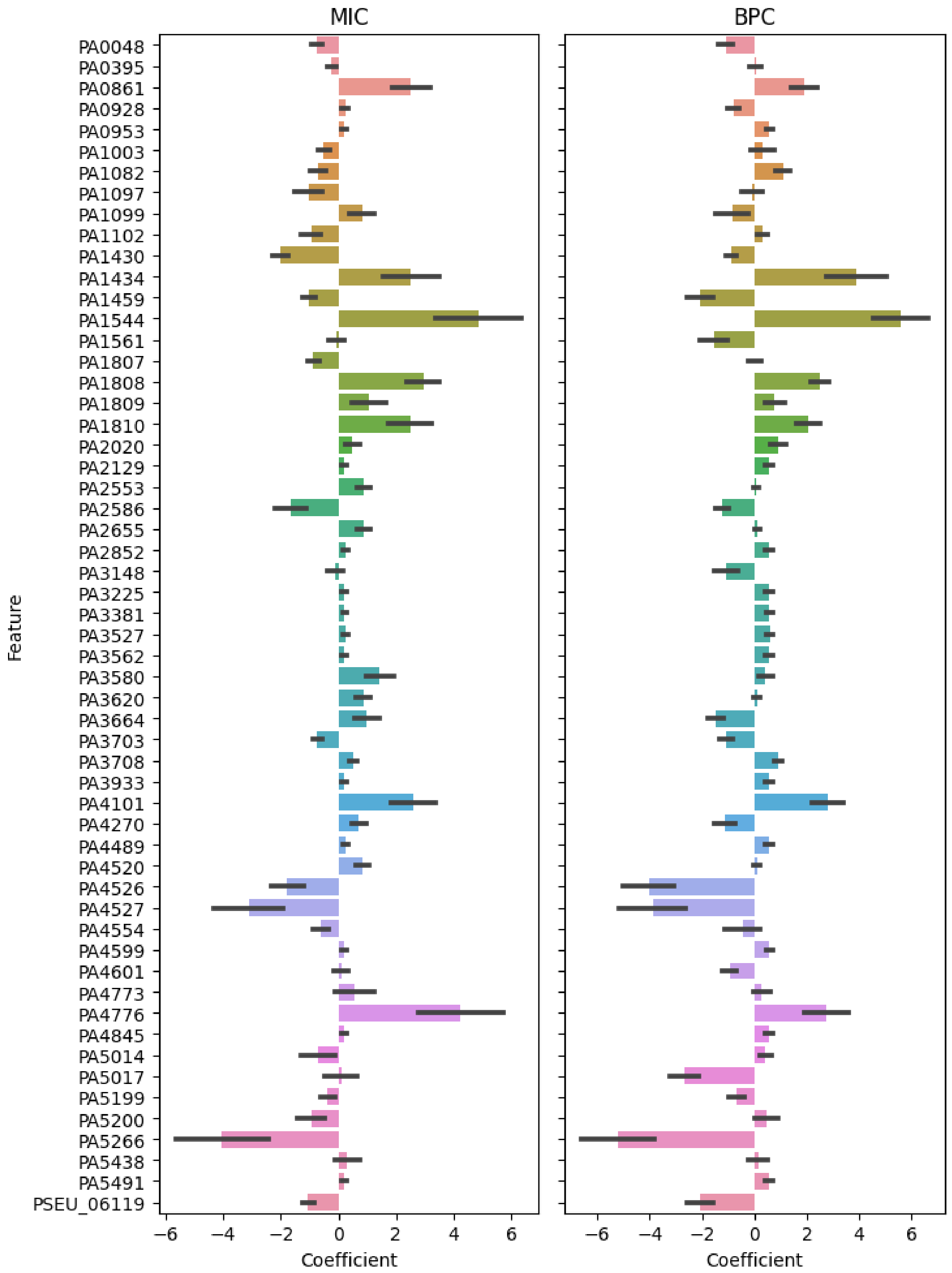


**Figure S7 Learned model coefficients for DNA variants**. Coefficients refer to the linear transformation coefficients $\beta$ in the ordinal regression model. The left-hand and right-hand sides show coefficients for the model trained to predict minimal inhibitory concentration (MIC) and biofilm prevention concentration (BPC), respectively.
