## Supplementary material for "Harnessing machine learning to predict antibiotic susceptibility in *Pseudomonas aeruginosa* biofilms": Supplemntal figure S3

### ***WGS_MIC_predictions***

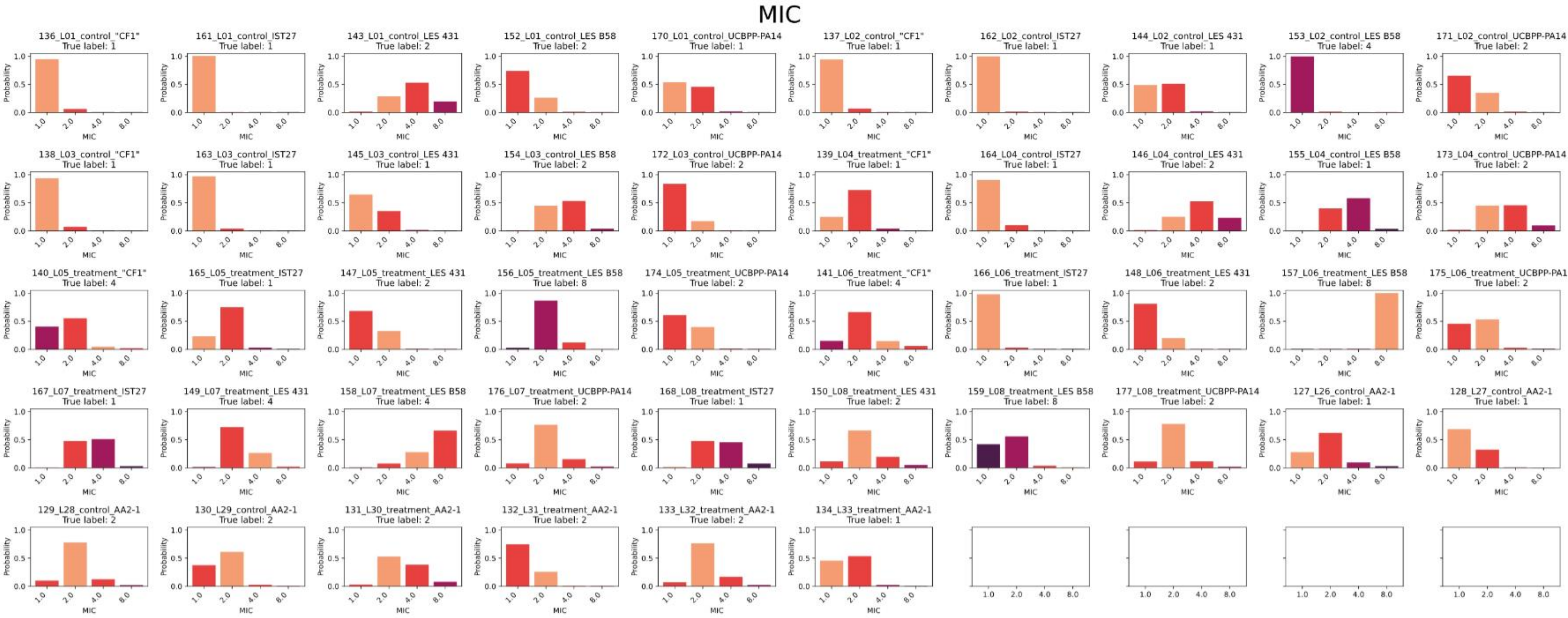

### *WGS_BPC_predictions*

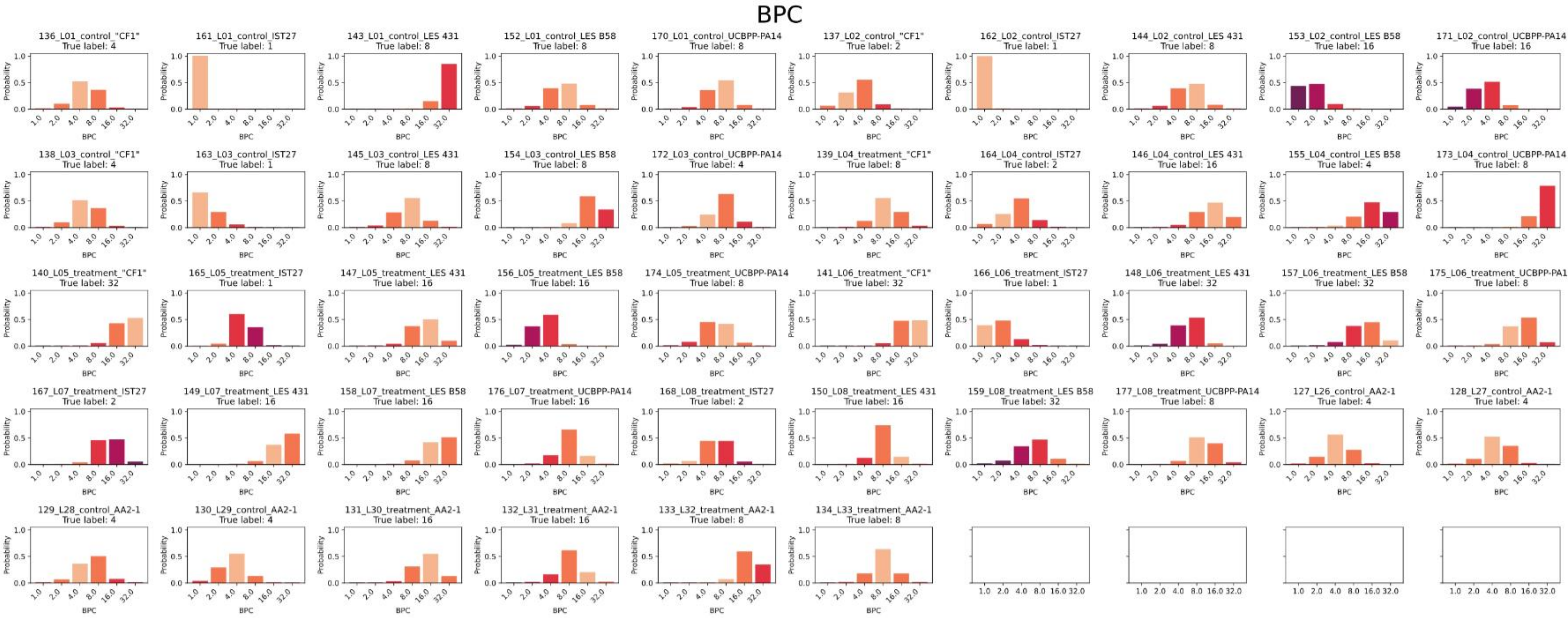

### *MALDI_MIC_predictions*

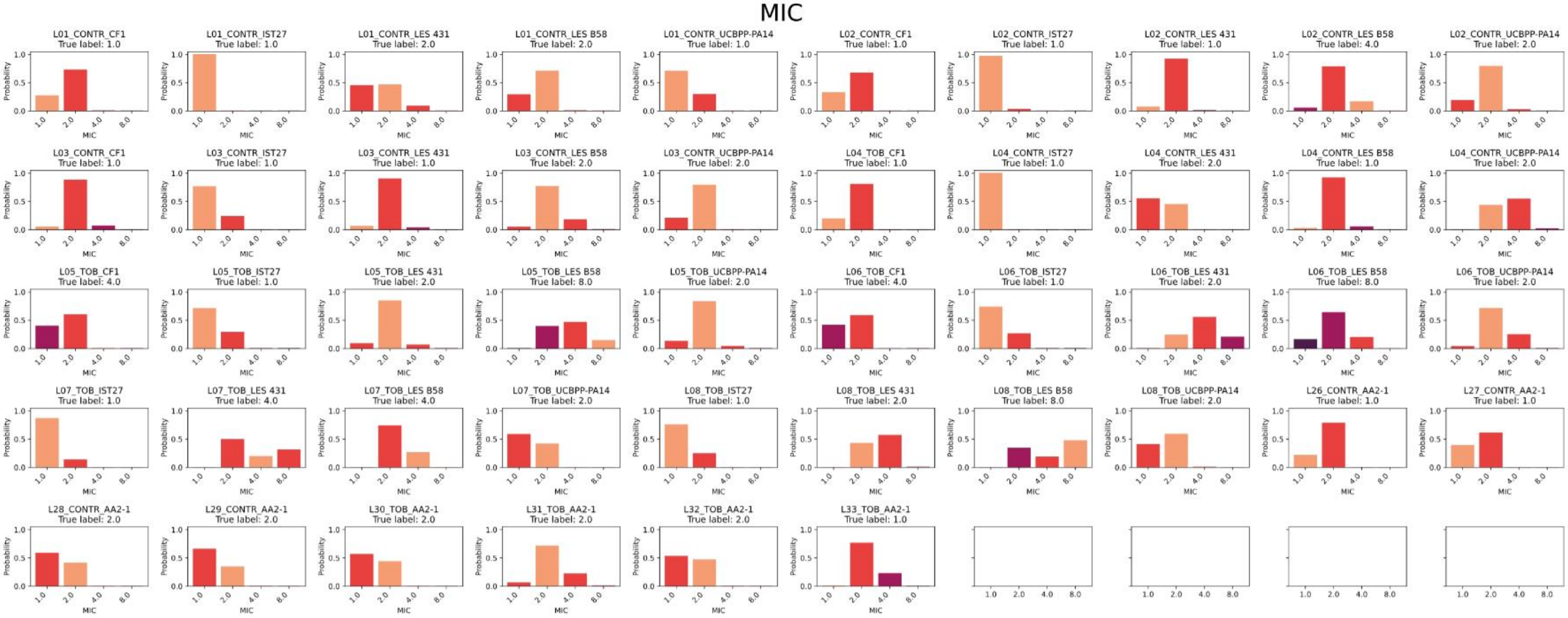

### *MALDI_BPC_predictions*

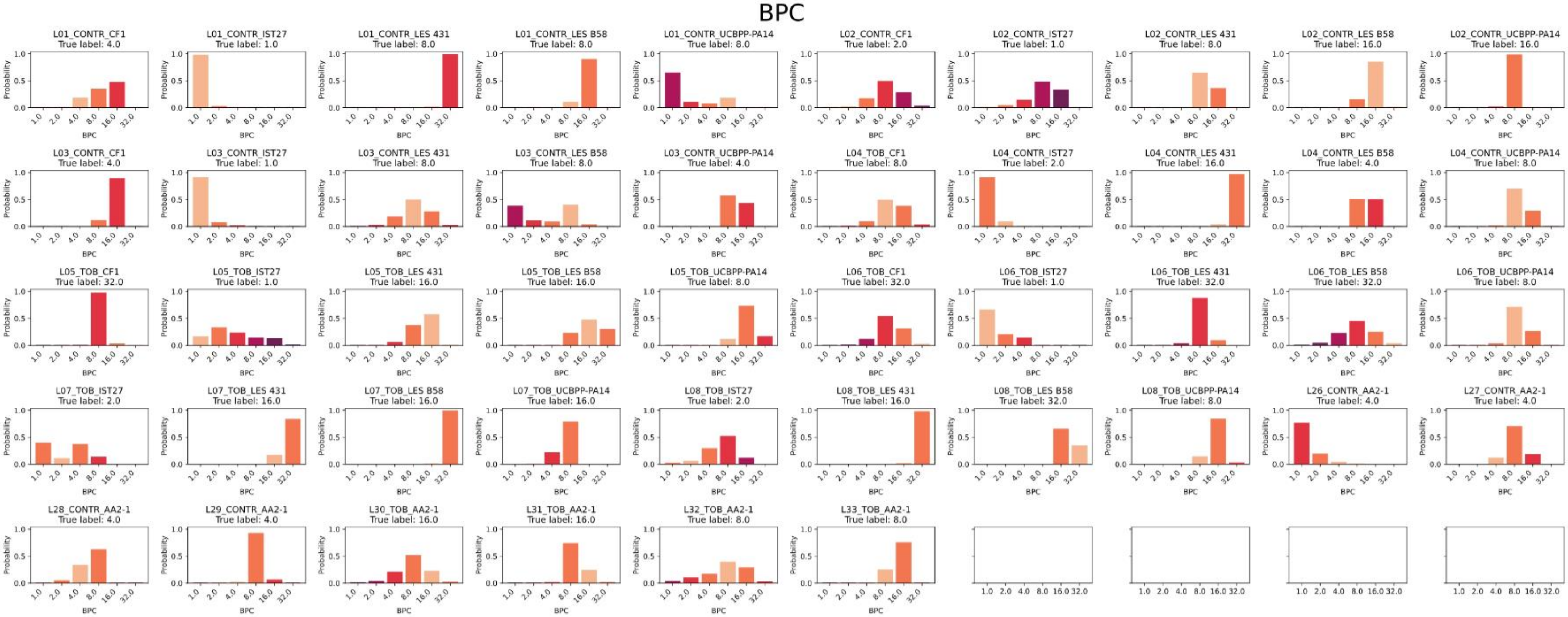

### *IMC_MIC_predictions*

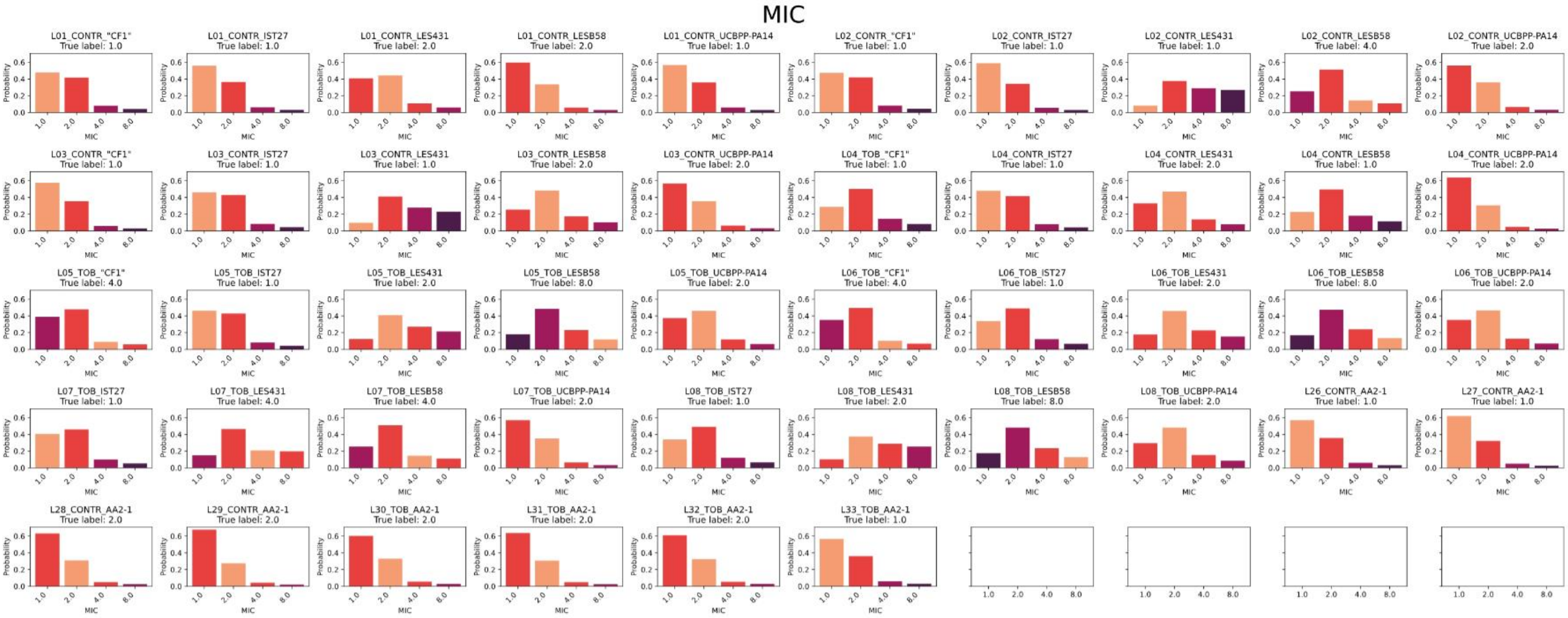

### *IMC_BPC_predictions*

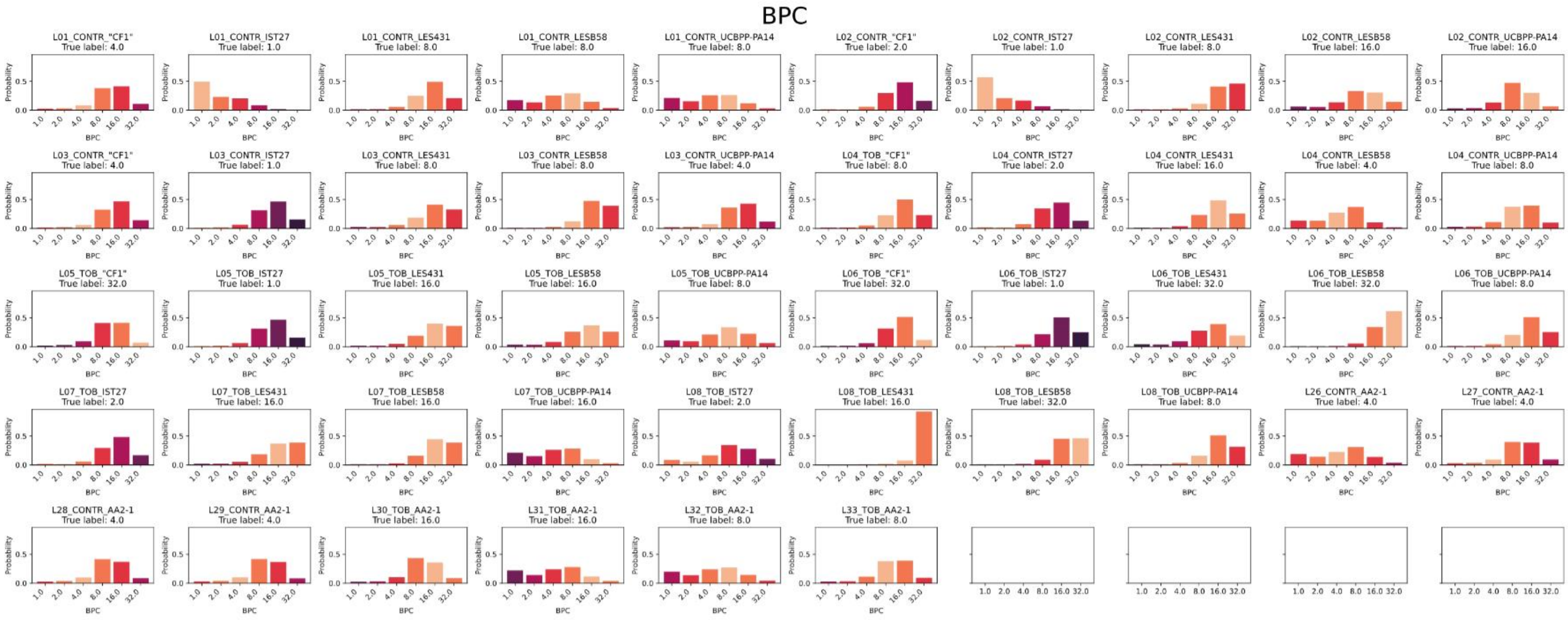

### *Raman_532_MIC_predictions*

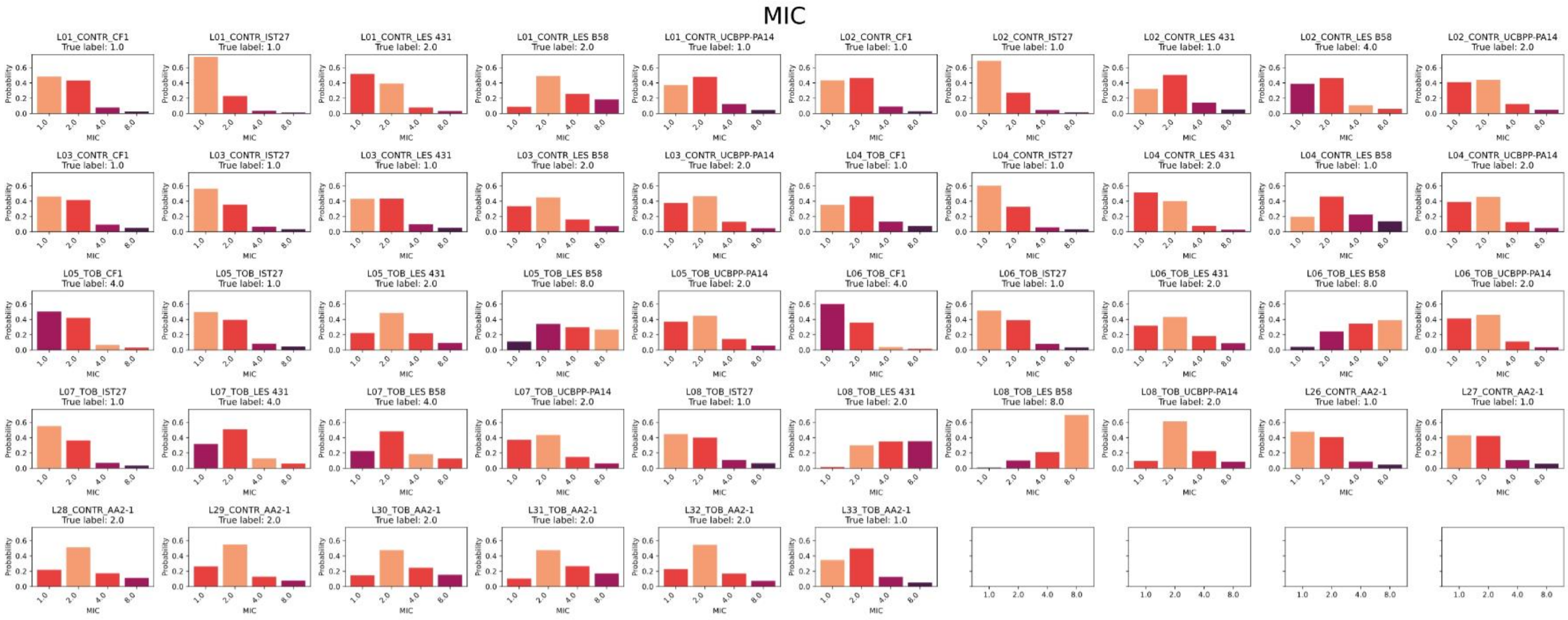

### *Raman_532_BPC_predictions*

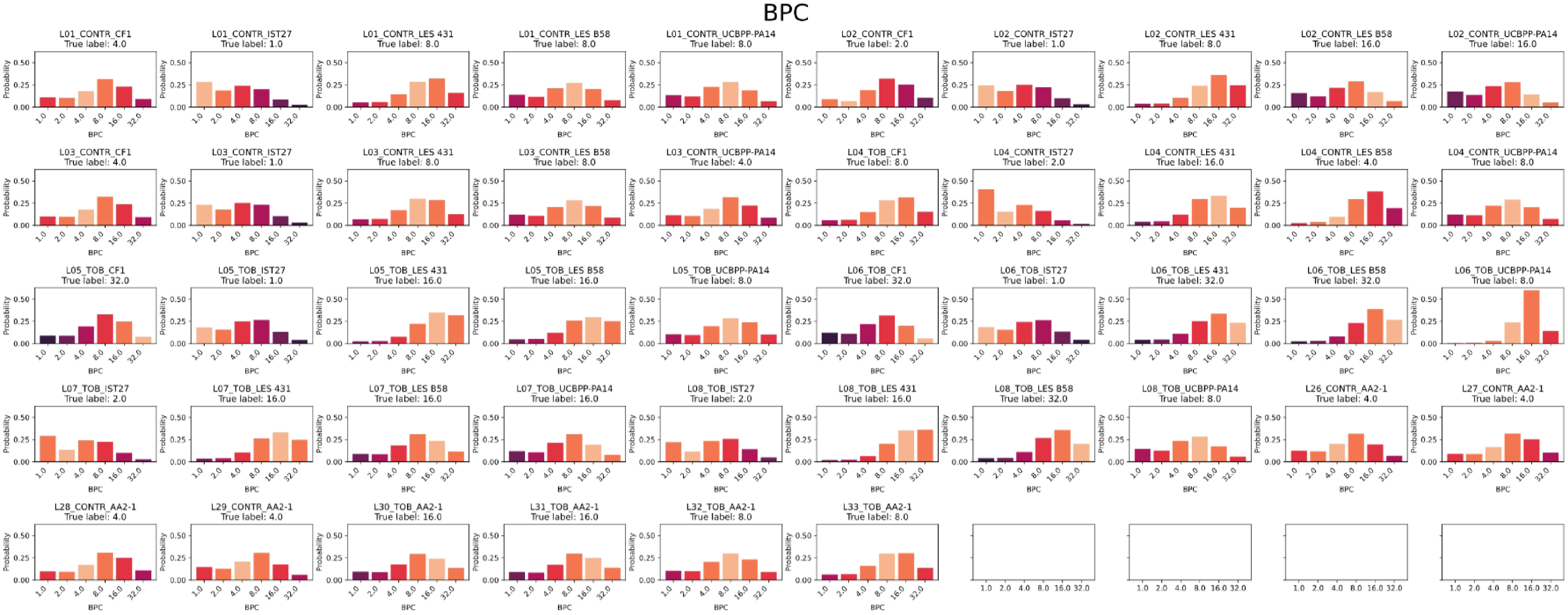

### *Raman_785_MIC_predictions*

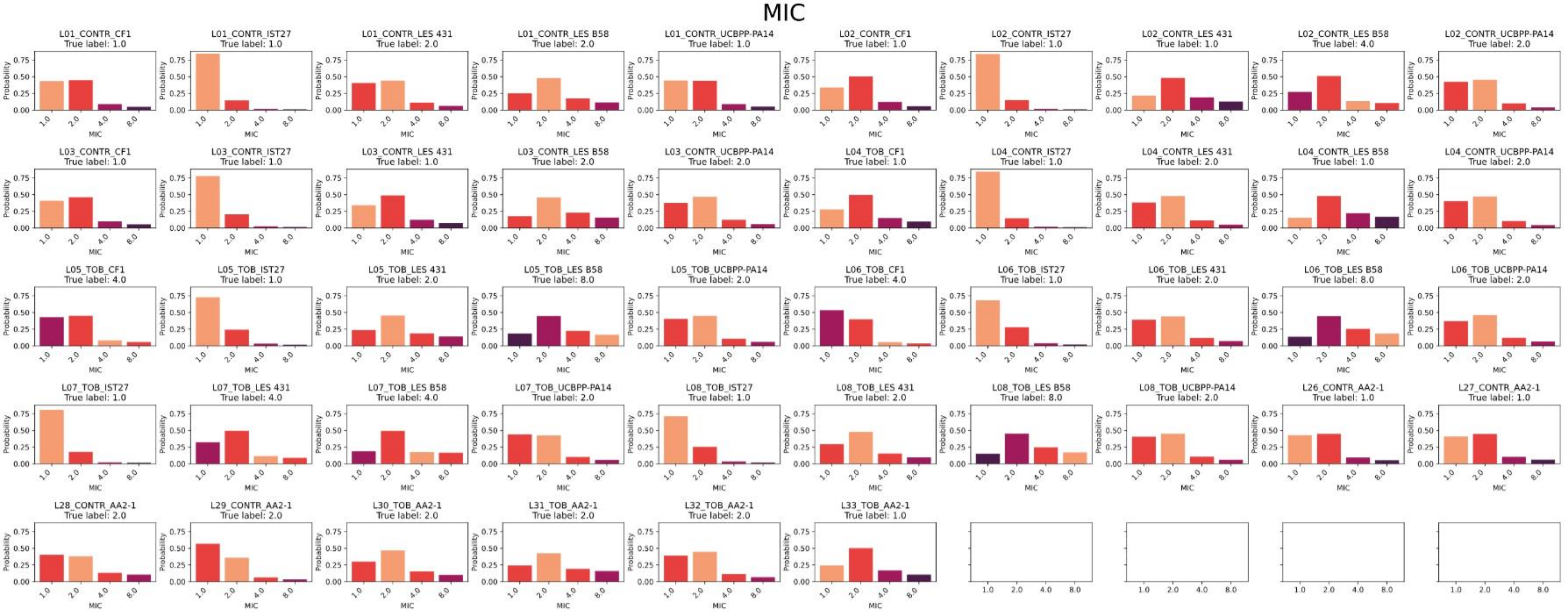

### *Raman_785_BPC_predictions*

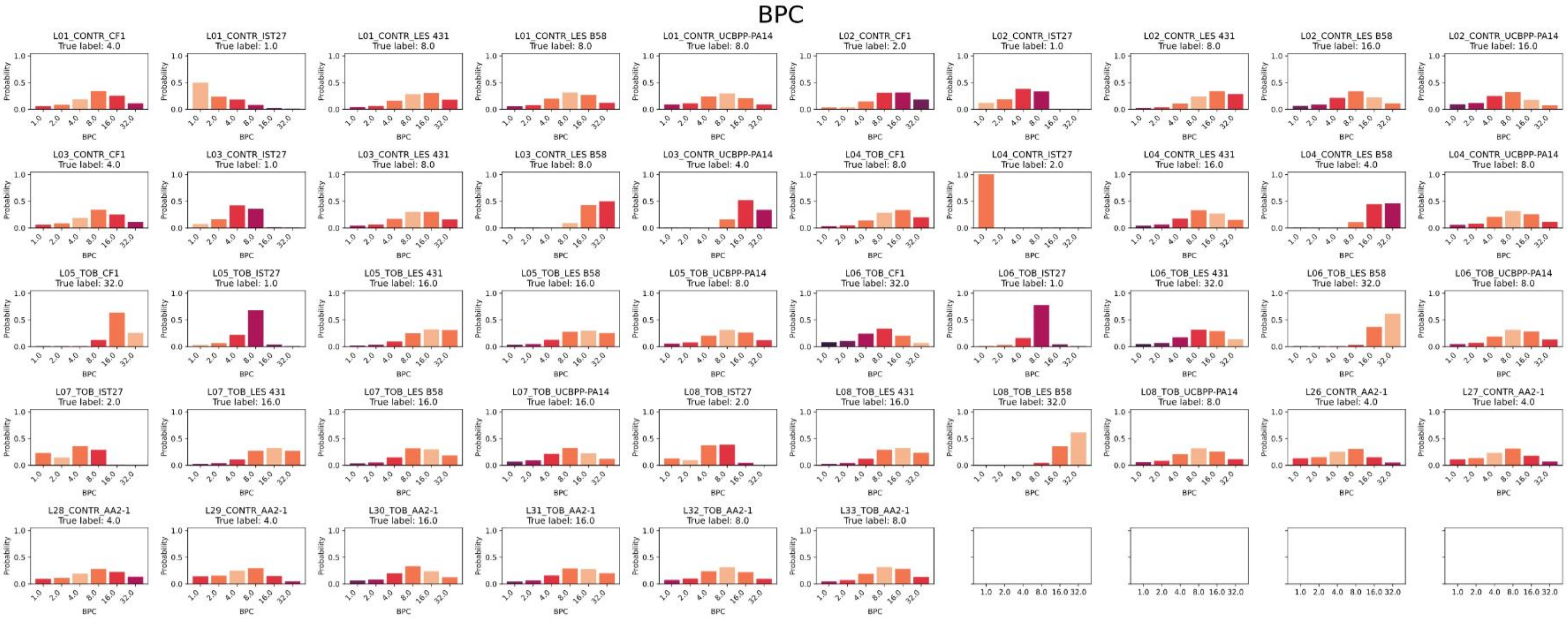

### *MX-Raman_MIC_predictions*

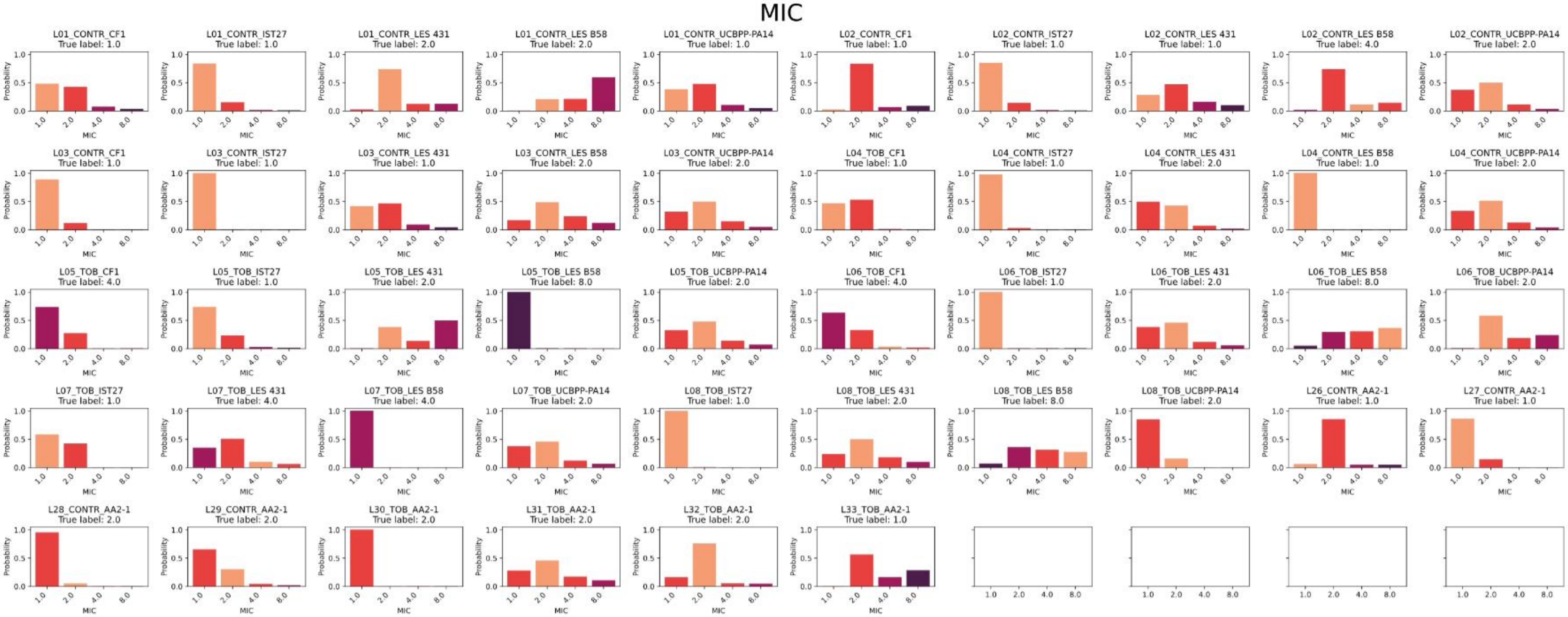

### *MX-Raman_BPC_predictions*

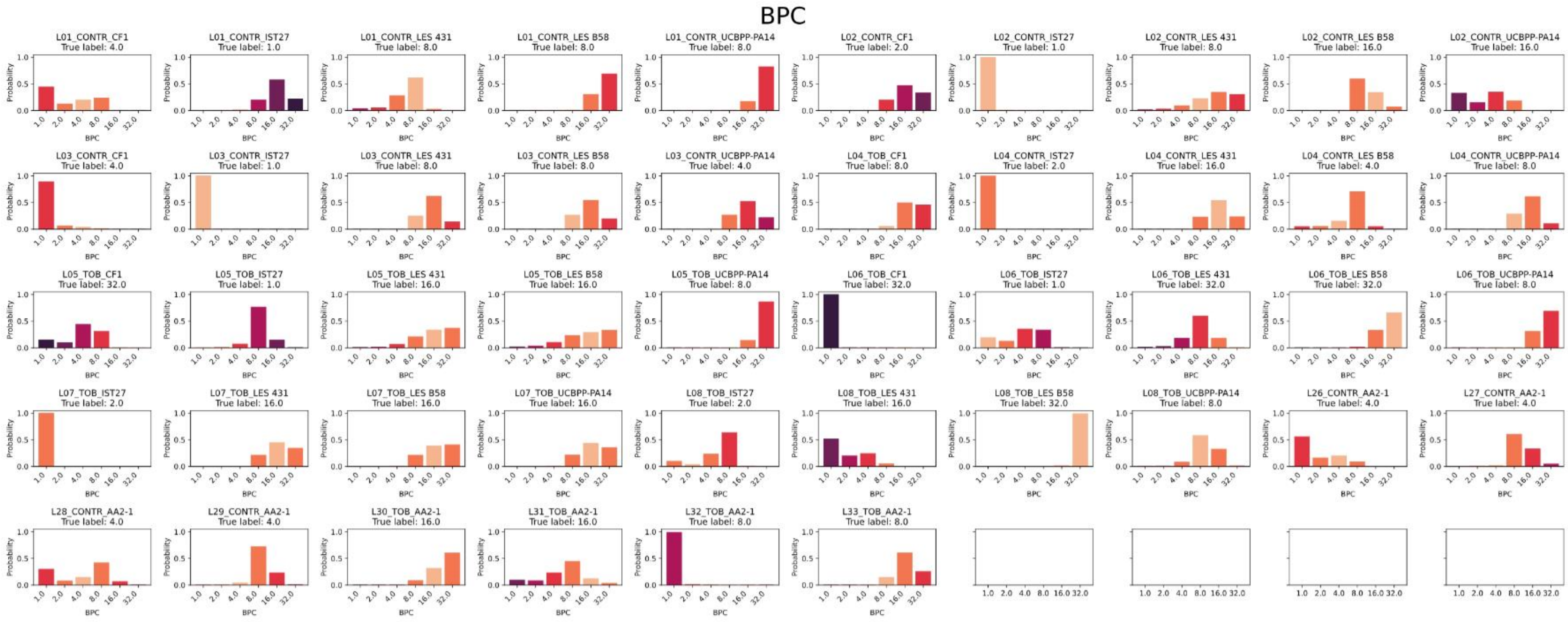
